# A common CTRB misfolding variant associated with pancreatic cancer risk causes ER stress and inflammation in mice

**DOI:** 10.1101/2024.07.23.604778

**Authors:** Cristina Bodas, Irene Felipe, Brice Chanez, Miguel Lafarga, Evangelina López de Maturana, Jaime Martínez-de-Villarreal, Natalia del Pozo, Marina Malumbres, Pierfrancesco Vargiu, Ana Cayuela, Isabel Peset, Katelyn E. Connelly, Jason W. Hoskins, Raúl Méndez, Laufey T. Amundadottir, Núria Malats, Sagrario Ortega, Francisco X. Real

## Abstract

**Objective:** Genome wide association studies have identified an exon 6 *CTRB2* deletion variant that associates with increased risk of pancreatic cancer. To acquire evidence on its causal role, we developed a new mouse strain carrying an equivalent variant in *Ctrb1*, the mouse orthologue of *CTRB2*.

**Design:** We used CRISPR/Cas9 to introduce a 707bp deletion in *Ctrb1* encompassing exon 6 (*Ctrb1*^Δexon6^). This mutation closely mimics the human deletion variant. Mice carrying the mutant allele were extensively profiled at 3 months to assess their phenotype.

**Results:** *Ctrb1*^Δexon6^ mutant mice express a truncated CTRB1 that accumulates in the ER. The pancreas of homozygous mutant mice displays reduced chymotrypsin activity and total protein synthesis. The histological aspect of the pancreas is inconspicuous but ultrastructural analysis shows evidence of dramatic ER stress and cytoplasmic and nuclear inclusions. Transcriptomic analyses of the pancreas of mutant mice reveals acinar program down-regulation and increased activity of ER stress-related and inflammatory pathways. Heterozygous mice have an intermediate phenotype. *Agr2* is one of the most up-regulated genes in mutant pancreata. *Ctrb1*^Δexon6^ mice exhibit impaired recovery from acute caerulein-induced pancreatitis. Administration of TUDCA or sulindac partially alleviates the phenotype. A transcriptomic signature derived from the mutant pancreata is significantly enriched in normal human pancreas of *CTRB2* exon 6 deletion variant carriers from the GTEx cohort.

**Conclusions:** This mouse strain provides formal evidence that the *Ctrb1*^Δexon6^ variant causes ER stress and inflammation *in vivo*, providing an excellent model to understand its contribution to pancreatic ductal adenocarcinoma development and to identify preventive strategies.

**SUMMARY BOX:** *What is already known about this subject?:*

- CTRB2 is one of the most abundant proteins produced by human pancreatic acinar cells.
- A common exon 6 deletion variant in *CTRB2* has been associated with an increased risk of pancreatic ductal adenocarcinoma.
- Misfolding of digestive enzymes is associated with pancreatic pathology.

*What are the new findings?:*

- We developed a novel genetic model that recapitulates the human *CTRB2* deletion variant in the mouse orthologue, *Ctrb1*.
- Truncated CTRB1 misfolds and accumulates in the ER; yet, mutant mice display a histologically normal pancreas at 3 months age.
- CTRB1 and associated chaperones colocalize in the ER, the cytoplasm, and the nucleus of acinar cells.
- Transcriptomics analysis reveals reduced activity of the acinar program and increased activity of pathways involved in ER stress, unfolded protein response, and inflammation.
- Mutant mice are sensitized to pancreatic damage and do not recover properly from a mild caerulein-induced pancreatitis.
- TUDCA administration partially relieves the ER stress in mutant mice.

*How might it impact on clinical practice in the foreseeable future?:*

- The new mouse model provides a tool to identify the mechanisms leading to increased pancreatic cancer risk in *CTRB2* exon 6 carriers.
- The findings suggest that drugs that cause ER stress relief and/or reduce inflammation might provide preventive opportunities.

## INTRODUCTION

Pancreatic Ductal Adenocarcinoma (PDAC) is the most common tumor type of the pancreas (>90%) and is predicted to become the second cause of cancer-related death in the Western world by 2030^1^. Both genetic and non-genetic factors contribute to PDAC risk ^2, 3^. Genome wide association studies (GWAS) have identified low-penetrance variants associated with an increased PDAC risk in the vicinity of genes related to acinar biology, including transcription factors (e.g., *NR5A2*, *HNF1A*), digestive enzymes (i.e., *CTRB1/2*) and endoplasmic reticulum (ER) stress factors (i.e., *XBP1*) ^4, 5^. Genetic variation in some of these genes has also been shown to be involved in susceptibility for chronic pancreatitis (CP), a well-established PDAC risk factor ^2, 3, 6^.

Genetically engineered mouse models (GEMMs) recapitulating PDAC initiation and progression are widely used. Expression of mutant *Kras* in the embryonic pancreas leads to PanINs and PDAC (reviewed in ^7, 8^). In contrast, mutant *Kras* activation in adult acinar cells is insufficient to cause PDAC but, when combined with acute or chronic pancreatitis, PanIN and PDAC develop ^7, 8,9^. These findings indicate that acinar cells can give rise to tumors with ductal features, placing them in the center of PDAC pathogenesis. Therefore, understanding acinar cell biology is key to unraveling how PDAC develops. We and others have shown that genetic alterations involving transcription factors required for acinar maturation sensitize acinar cells to the oncogenic effects of mutant *Kras*^10–15^.

Pancreatic acinar cells are highly specialized protein factories synthesizing massive amounts of hydrolytic enzymes. Acinar cells have a well-developed rough ER, a prominent Golgi complex, and abundant secretory granules, and they show constitutive low-level ER stress. Inappropriate intrapancreatic activation of digestive enzymes plays a crucial role in acute and chronic pancreatitis^16^. Rare germline mutations in several genes coding for digestive enzymes (e.g., *PRSS1, CPA1*) have been shown to cause hereditary CP and increase the risk of PDAC ^17–19^. However, these mutations are very rare in PDAC patients.

Recently, GWAS have identified common germline variants in the *CTRB1/2* locus at human chromosome 16q23.1 as risk factors for CP ^20^ and sporadic PDAC ^5, 21^. Variants in this locus have also been associated with the risk of type 1 and type 2 diabetes mellitus ^5^. Fine mapping of the PDAC GWAS signals led to the identification of a *CTRB2* exon 6 deletion, present in 16% of the population in heterozygosity and in 0.6% in homozygosity, as a putative causal variant associated with increased PDAC risk ^21^. *CTRB2* encodes chymotrypsinogen B2, one of the most abundant proteins produced by acinar cells. Ectopic expression of a *CTRB2* cDNA carrying the exon 6 deletion in cultured cells results in a truncated protein that misfolds and accumulates in the ER, inducing ER stress ^21^. However, formal evidence for a causal role of the deletion is missing and will be hard to obtain in humans.

Chymotrypsin plays a key role in pancreatitis: *Ctrb1* knockout mice are normal up to one year of age but they display increased intrapancreatic trypsin activity and a more severe pancreatitis upon caerulein administration ^22^. These observations support the hypothesis that chymotrypsin limits the activity of trypsin through its degradation. Here, we have used CRISPR/Cas9 to establish a new mouse strain carrying the equivalent human exon 6 deletion in *Ctrb1*, the only gene coding for chymotrypsinogen in mice (*Ctrb1*^Δexon6^). We have extensively phenotyped 3 month-old mutant mice. Using a combination of *in vivo* studies, ultrastructural and omics analyses we uncover that the exon 6 deletion causes massive ER stress, down-regulation of the acinar differentiation program, inflammation, and DNA damage. Preliminary data indicate that this phenotype may be partially relieved by chaperone-like drugs. The transcriptomic features of the mutant mice recapitulate features of the normal pancreas of carriers of the *CTRB2* exon 6 deletion variant from the GTEx cohort.

## MATERIALS AND METHODS

### CRISPR reagents

CRISPR gRNAs were designed using the CRISPOR web tool (crispor.tefor.net) to target sequences in introns 5 and 6 of mouse *Ctrb1-201*, chromosome 8, reverse strand (www.ensembl.org). All CRISPR reagents were purchased from Integrated DNA Technologies Inc. (IDT). gRNAs were assembled from Alt-R® CRISPR-Cas9 crRNAs and Alt-R® CRISPR-Cas9 tracrRNA. The two crRNAs containing the sequences 5′TTGTTAATTAAGGCTGGTCG3′ (forward strand, targeting intron 5) and 5′CTAGCCGGATCAGGTTAGGC 3′ (reverse strand, targeting intron 6) were annealed to the corresponding TracRNAs in microinjection buffer (10 mM Tris/HCl pH 7,5; 0.1 mM EDTA) by heating 5 min at 95 °C in a thermocycler and decreasing temperature slowly using a pre-defined temperature ramp. Assembled Trac/Cr gRNAS were incubated with Alt-R S.p. Cas9 Nuclease V3 to form the ribonucleoprotein complex 15 min at room temperature in microinjection buffer.

### Zygote microinjection and line generation

CRISPR reagents were microinjected in the pronucleus of zygotes obtained from crosses between hybrid B6.CBA males and females. Females (5-8 week-old) were previously superovulated by consecutive administration of 5 IU of PMSG (at 3 pm of day -3) and 5 IU of hCG (at 1 pm of day -1) and matings were set up immediately after hCG administration. At 8 pm of day 0, the formation of vaginal plugs was monitored and cumuli were collected from oviducts. Cumuli were disaggregated with a hyaluronidase (Sigma H4272) solution (10 mg/mL in M2 medium) and diluted 1:2 in M2 medium at treatment. Free zygotes were cultured in KSOM medium until injection. Concentrations of CRISPR reagents for microinjection were 2.5 mM Trac/Cr for each gRNA and 0.24 mM Cas9 in microinjection buffer. Next day, zygotes that had developed to 2-cell stage were transferred to pseudopregnant CD1 females according to standard protocols ^23^. F0 pups were initially genotyped by PCR using the following primers: GGTTGTGAACCACCATGTGG (forward) and GACAAGGATACCCAGA (reverse). Expected amplicons were 1079 bp (WT) and 420 bp (Δexon6). Out of 35 pups born, 15 had the expected Δexon6 band. The Δexon6 amplicon was sequenced in 6 of the positive pups with the primers used for PCR amplification and the correct deletion edges were confirmed. Two pups were chosen as founders and crossed with C57BL6 females to establish two independent lines. The same analysis was repeated in the F1; one F1 animal from each line was considered founder. Wild type (WT) and homozygous mice yield amplicons of 670bp and 374bp, respectively. Heterozygous mice display both amplicons.

### Statistical analysis

All group data are represented by the mean; error bars are the standard deviation (SD). Student’s t-test was used to assess the difference in means between two groups of a normally-distributed continuous variable. For comparisons not showing a normal distribution, two-tailed Mann–Whitney U-test was used. Two-tailed p-values<0.05 were considered significant. All statistical analyses were performed with GraphPad Prism v8. The statistical test used in each experiment is indicated in the Figure legends or in the Source Data file.

Detailed methods are provided in the **Supplemental Materials** file.

## RESULTS

### Generation and validation of a *Ctrb1* exon 6 deletion mutant mouse model

To acquire formal evidence on the effects of the *CTRB2* exon 6 deletion on pancreatic homeostasis, we developed a novel mouse model recapitulating the human variant in the mouse ortholog, *Ctrb1*. Single cell RNA-Seq data from the Tabula Muris Consortium indicate that *Ctrb1* mRNA is expressed only in the pancreas (**Supplemental Figure 1A**) and the protein is detected in embryonic pancreas at e14.5 (**Supplemental Figure 1B**). Mouse CTRB1 and human CTRB2 display a 93% protein homology (**Supplemental Figure 1C**) and their transcripts are among the highest in the adult pancreas. *CTRB1* and *CTRB2* transcripts are expressed at comparable levels in human pancreas (**Supplemental Table 1**). **Figure 1A** shows the strategy used to generate an exon 6 deletion in *Ctrb1* that mirrors the human *CTRB2* variant. Wild type CTRB2 has 263 amino acids; the 584 bp deletion is predicted to yield a truncated 166 residue protein due to a premature stop codon in exon 7. The corresponding 659bp exon 6 deletion in *Ctrb1* is predicted to produce a truncated protein of the same length (**Supplemental Figure 1C**).

**Figure 1.**
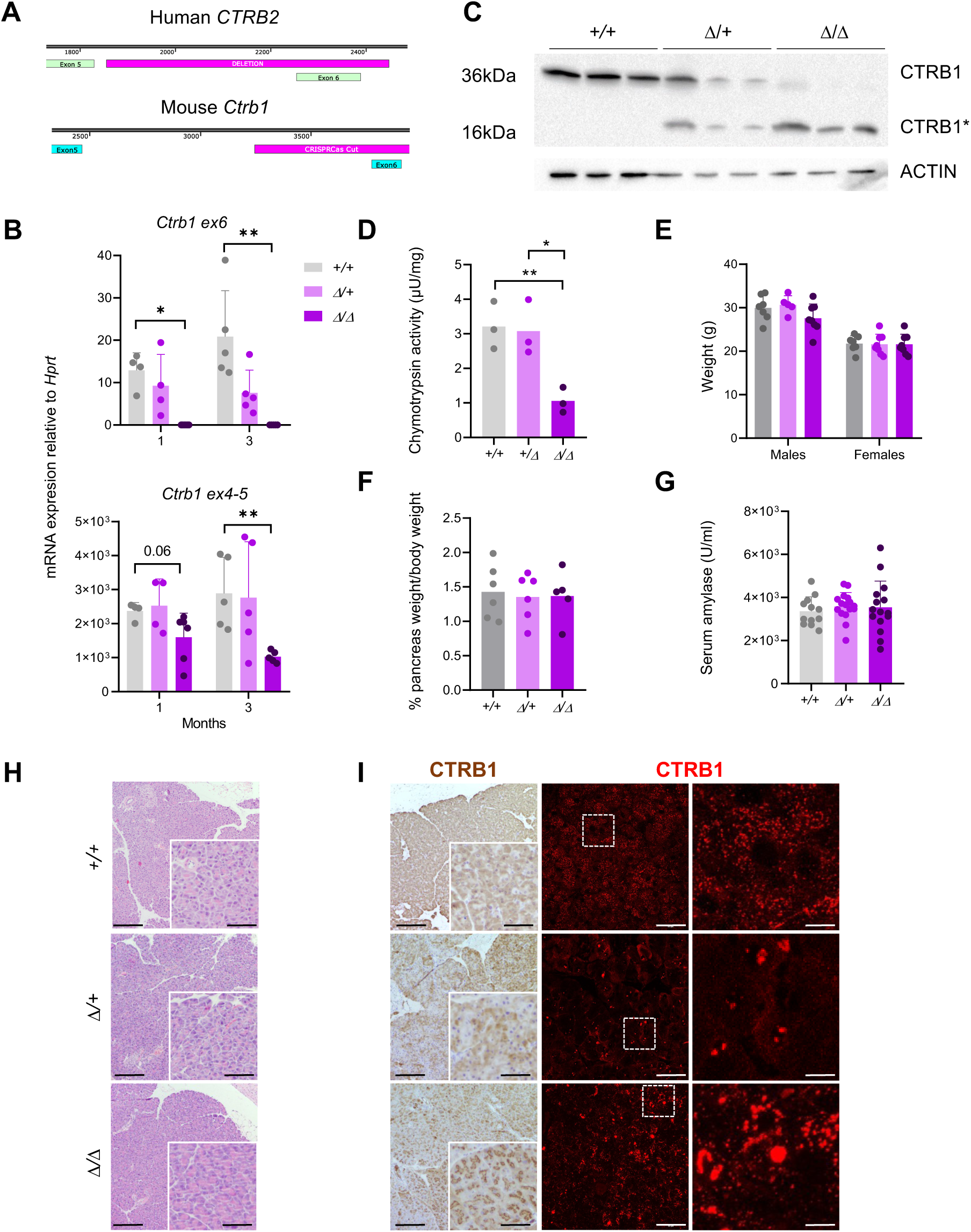
*Ctrb1^△ex6^* mice recapitulate human *CTRB2* exon 6 deletion and express a truncated CTRB1 protein that forms aggregates. **A**. Schematic representation of the human and mouse exon 6 deletion. **B**. *Ctrb1* exon 6 sequences are undetectable, and *Ctrb1* exon 4-5-containing transcripts levels are lower, in *Ctrb1^Δ/Δ^* mice (n≥4/group). **C**. Mice carrying the *Ctrb1* exon 6 deletion allele express a smaller, truncated, protein (n=3/group). **D**. Chymotrypsin activity is reduced in *Ctrb1^Δ/Δ^* mice (n=3/group). **E-G**. *Ctrb1^Δ/Δ^* mice show normal body weight (n≥6/group) (**E**), pancreas weight/body weight ratio (n≥5/group) (**F**), and normal serum amylase levels (n≥12/group) (**G**). **H.** *Ctrb1*^Δ/Δ^ pancreata display normal histology. Scale bar, 200µm; inset scale bar, 50µm. **I**. CTRB1 accumulates as aggregates in *Ctrb1^Δ/Δ^* pancreata. Representative images are shown (from n≥6 mice). Scale bar IHC, 200µm; inset scale bar, 50µm. Scale bar IF, 50µm; inset scale bar, 5µm.

The genotyping strategy used is shown in **Supplemental Figure 1D**. **Figure 1B** shows that exon 6-containing transcripts were reduced by approximately 50% in 1- and 3-month-old *Ctrb1^+/Δ^* mice and were undetectable in *Ctrb1^Δ/Δ^* mice. Total *Ctrb1* mRNA expression levels - quantified using primers spanning exons 4-5 - were significantly lower in *Ctrb1^Δ/Δ^* mice (-3.2-fold, p-value<0.01). Anti-CTRB antibodies detected a 36 kDa protein in pancreatic lysates from WT mice, whereas a truncated protein of approximately 16 kDa was detected in *Ctrb1^Δ/Δ^* pancreata. Both protein species were detected in lysates from *Ctrb1^+/Δ^* mice (**Figure 1C**). Chymotrypsin activity was significantly reduced in homozygous mutant mice (**Figure 1D**). These results validate the predicted synthesis of a truncated CTRB protein from the mutant allele.

There were no significant differences in body weight or pancreas/body weight of 1- and 3-month-old WT, *Ctrb1^+/Δ^*, and *Ctrb1^Δ/Δ^* mice (**Figure 1E,F**). Baseline serum amylase levels were also similar (**Figure 1G**). Histological analysis did not reveal major alterations in the pancreas of *Ctrb1^+/Δ^* and *Ctrb1^Δ/Δ^* mice (**Figure 1H**). However, immunohistochemistry (IHC) and immunofluorescence (IF) revealed striking differences in CTRB1 expression: a fine punctate pattern was observed in WT acinar cells whereas a weaker expression, associated with coarse speckles suggestive of protein aggregates, was noted in *Ctrb1^+/Δ^* and *Ctrb1^Δ/Δ^* pancreata (**Figure 1I**).

### Truncated CTRB1 forms cytosolic and nuclear aggregates

To determine whether the endogenous mutant protein accumulates in the ER, we stained pancreatic sections with antibodies against CTRB1 and CALR (a rough ER marker) (**Figure 2A**). CALR immunostaining showed a higher signal intensity in *Ctrb1^Δ/Δ^* than in WT pancreata. At 1 month, CALR and CTRB1 co-localized as discrete dots in acinar cells of homozygous mutant mice; however, at 3 months CTRB1 showed a coarser pattern with prominent cytoplasmic aggregates or inclusions lacking CALR. The number of cytosolic aggregates was higher in *Ctrb1^Δ/Δ^* acinar cells than in WT cells. Significantly more cytoplasmic aggregates were seen in mice at 1 month compared to 3 months of age (**Figure 2B**), although they were larger at 3 months (**Figure 2A,C**). CTRB1 aggregates were also localized in the nucleus of acinar cells in *Ctrb1^Δ/Δ^* mice, but not in WT mice. Using Proteostat^®^, a significantly higher amount of misfolded proteins was detected in *Ctrb1^Δ/Δ^* than in WT pancreata (53% higher, p-value=0.028) (**Figure 2D**). Mutant CTRB1 was found both in the SDS-soluble and - insoluble fractions of lysates, whereas wild type CTRB1 was found almost exclusively in the SDS-soluble fraction (**Figure 2E**). We also found ER stress-related chaperones, such as BiP, in both fractions in *Ctrb1^Δ/+^* and *Ctrb1^Δ/Δ^* lysates. In addition, CTRB1 co-localized with BiP in the nucleus (**Figure 2F**).

**Figure 2.**
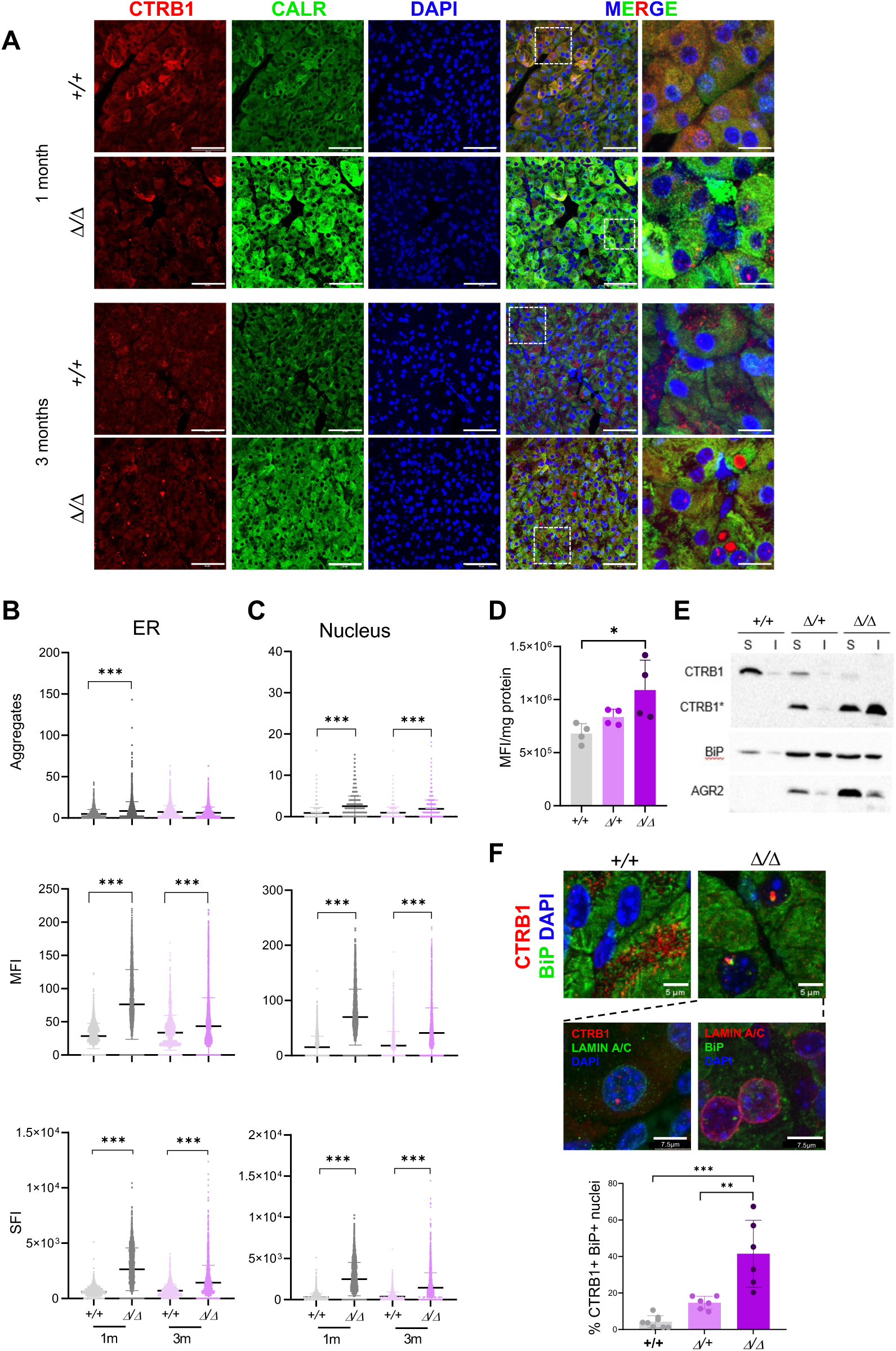
CTRB1 forms insoluble cytoplasmic and nuclear aggregates in *Ctrb1^Δ/Δ^* pancreata. **A**. Representative images showing CTRB1 aggregates and higher expression of calreticulin (CALR) in the pancreas of *Ctrb1^Δ/Δ^* mice. Scale bar, 50µm; inset scale bar, 5µm. **B, C**. Quantification of cytoplasmic (**B**) and nuclear (**C**) CTRB1 aggregates in acinar cells of *Ctrb1^Δ/Δ^*mice (n=3 mice/group, 5 images/case). **D**. The pancreas of *Ctrb1^Δ/Δ^* mice contains significantly more Proteostat^®^+ aggregates than that of wild type mice (n=3 mice/group). **E**. CTRB1 and AGR2 accumulate in the insoluble fraction only in *Ctrb1^Δ/+^* and *Ctrb1^Δ/Δ^* pancreata. BiP preferentially partitions to the insoluble protein fraction in *Ctrb1^Δ/+^* and *Ctrb1^Δ/Δ^* mice (n=3 mice/group). **F**. CTRB1 is co-expressed with BiP in the nucleus of *Ctrb1^Δ/Δ^* acinar cells (n≥6 mice/group). Nuclear localization is confirmed through double-labeling with lamin A/C.

### *Ctrb1^Δ/Δ^* pancreata display massive ER cisternal dilation and cytoplasmic and nuclear inclusions

The lack of overt histological abnormalities, despite the occurrence of CTRB aggregates, prompted a detailed ultrastructural analysis. The main findings are summarized in **Table 1**. Acinar cells from WT mice showed the expected morphology with abundant electron-dense zymogen granules and well-organized stacks of rough ER cisterna (**Figure 3A**). In contrast, acinar cells of *Ctrb1^Δ/Δ^*mice displayed a massive dilation of ER cisternae, accompanied by focal dilations of the perinuclear space at the nuclear envelope. In the lumen of ER cisternae, secretory products (e.g., truncated misfolded proteins) often coalesced to form larger aggregates or cytoplasmic inclusions, likely reflecting liquid-liquid phase separation forming intracisternal granules of biomolecular condensates (**Figure 3B,C**). Homozygous mutant mice also displayed an abnormally variable number and size of zymogen granules (**Figure 3D**). A subset of granules showed an electron-lucent peripheral halo that was absent from WT acinar cells (**Figure 3D**). A striking ultrastructural observation was the presence of nuclear inclusions in *Ctrb1^Δ/Δ^* acinar cells but not in WT cells (**Figure 3E**). A detailed description of the nuclear findings is provided in **Supplemental Figure 2**. In *Ctrb1^Δ/Δ^*pancreata, the interstitial tissue showed a normal organization, including capillaries and intercalated excretory ducts, with a few fibrocytes and isolated macrophages; inflammatory cells were inconspicuous **(Figure 3F**). Despite the germline nature of the *Ctrb1* mutation, acinar cells with an almost normal ultrastructural aspect were interspersed with the highly abnormal cells described above, indicating the occurrence of a mosaic phenotype.

**Figure 3.**
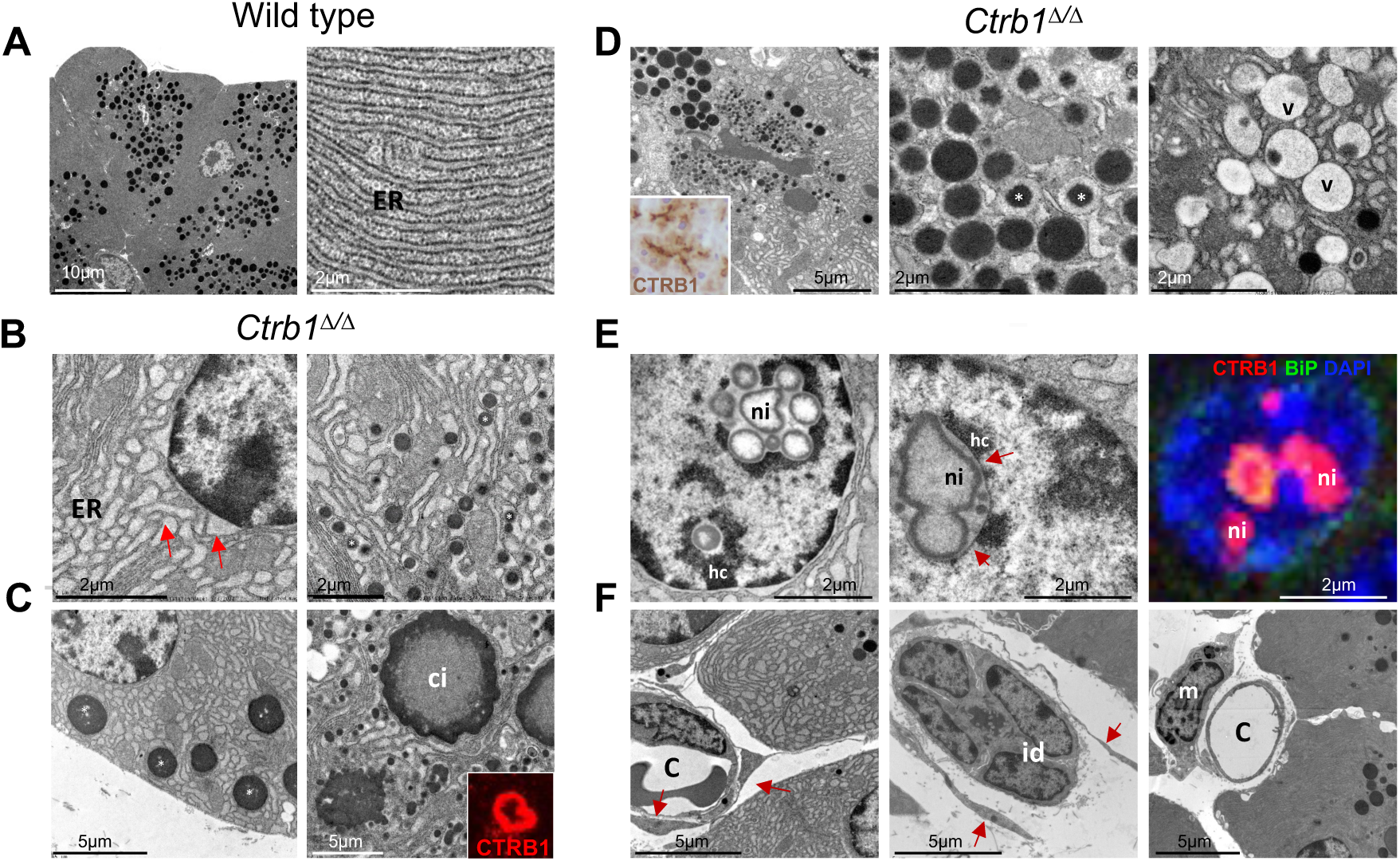
Transmission electron microscopy shows that acinar cells from *Ctrb1^Δ/Δ^* mice display dramatic ER stress and nuclear inclusions. **A**. Ultrastructural images showing normal acinar cell architecture in wild type pancreata. **B**. *Ctrb1^Δ/Δ^* acinar cells display dilation of the rough ER cisternae with focal dilations of the perinuclear space (red arrowheads) and intracisternal ER granules (right, asterisk). **C**. Coalescence of intracisternal granules in *Ctrb1^Δ/Δ^*pancreata, forming large aggregates (asterisks) or cytoplasmic inclusions (ci), likely corresponding to the coarse speckles observed using CTRB IF (right). Inset: cytoplasmic CTRB1+ cytoplasmic inclusion. **D**. Abnormal zymogen granule size heterogeneity (left) with corresponding CTRB-positive secretory products detected by IHC (inset). Cytoplasmic areas showing zymogen granules with an electron-lucent halo, suggestive of active proteolysis, surrounding a dense core (asterisks). Hydropic/vacuolar degeneration of zymogen granules (v). **E**. Nuclear inclusions (ni), likely representing the ultrastructural correlate of the nuclear CTRB1 detected using IF; nuclear CTRB1 (red) and BiP (green). Nuclear inclusions are surrounded by the inner nuclear membrane (arrows) and, partially, by heterochromatin (hc). **F**. Intralobular interstitial tissue with normal ultrastructural appearance, including capillaries (c) intercalated excretory ducts (id), scattered fibrocytes (red arrows) and isolated macrophages (m). Representative images from n=2 mice/group analyzed.

**Table 1.** Summary of the main ultrastructural findings in acinar cells.

| Age | 1 month | 3 months |
| --- | --- | --- |
| **Nucleus** |  |  |
| Intranuclear compact granules | - | ++ |
| Intranuclear hollow granules | - | ++ |
| Nuclear inclusions | - | + |
| Nucleolar alterations | - | + |
| Perinuclear cistern dilation | - | ++ |
| Heterochromatinization | - | ++ |
| **Cytoplasm** |  |  |
| ER dilation | - | +++ |
| ER disruption | - | + |
| Intracisternal granules | - | ++ |
| Intracisternal A-type retroviral particles | - | - |
| Cytoplasmic inclusions | - | ++ |
| Lipid droplets | - | - |
| Zymogen granule heterogeneity | + | ++ |
| Enlargement lysosomal compartment | - | - |
| Activation of autophagy | - | - |
| Ribophagy | - | - |
| Stress granules | - | - |
| Cellular degeneration | - | + |

### Transcriptomic analysis of the pancreas of *Ctrb1^Δ^* mutant mice

To gain a comprehensive understanding of the mechanisms underlying the phenotype of the pancreas of 3-month-old *Ctrb1^Δ/Δ^* mice, we performed genome wide transcriptomic analysis using RNA-seq. We observed 538 significantly Differentially Expressed Genes (DEGs), 291 of which were up-regulated and 247 down-regulated in *Ctrb1^Δ/Δ^* pancreata (**Figure 4A and Supplemental Table 2**). Pathway enrichment analysis using GSEA revealed a dramatic down-regulation of the acinar differentiation program and an up-regulation of ER stress- and inflammation-related pathways in *Ctrb1^Δ/Δ^* pancreata, with a milder, intermediate transcriptomic phenotype in *Ctrb1^+/Δ^* pancreata. Main findings are described below and shown in **Figures 4** and **Supplemental Figure 3**.

**Figure 4.**
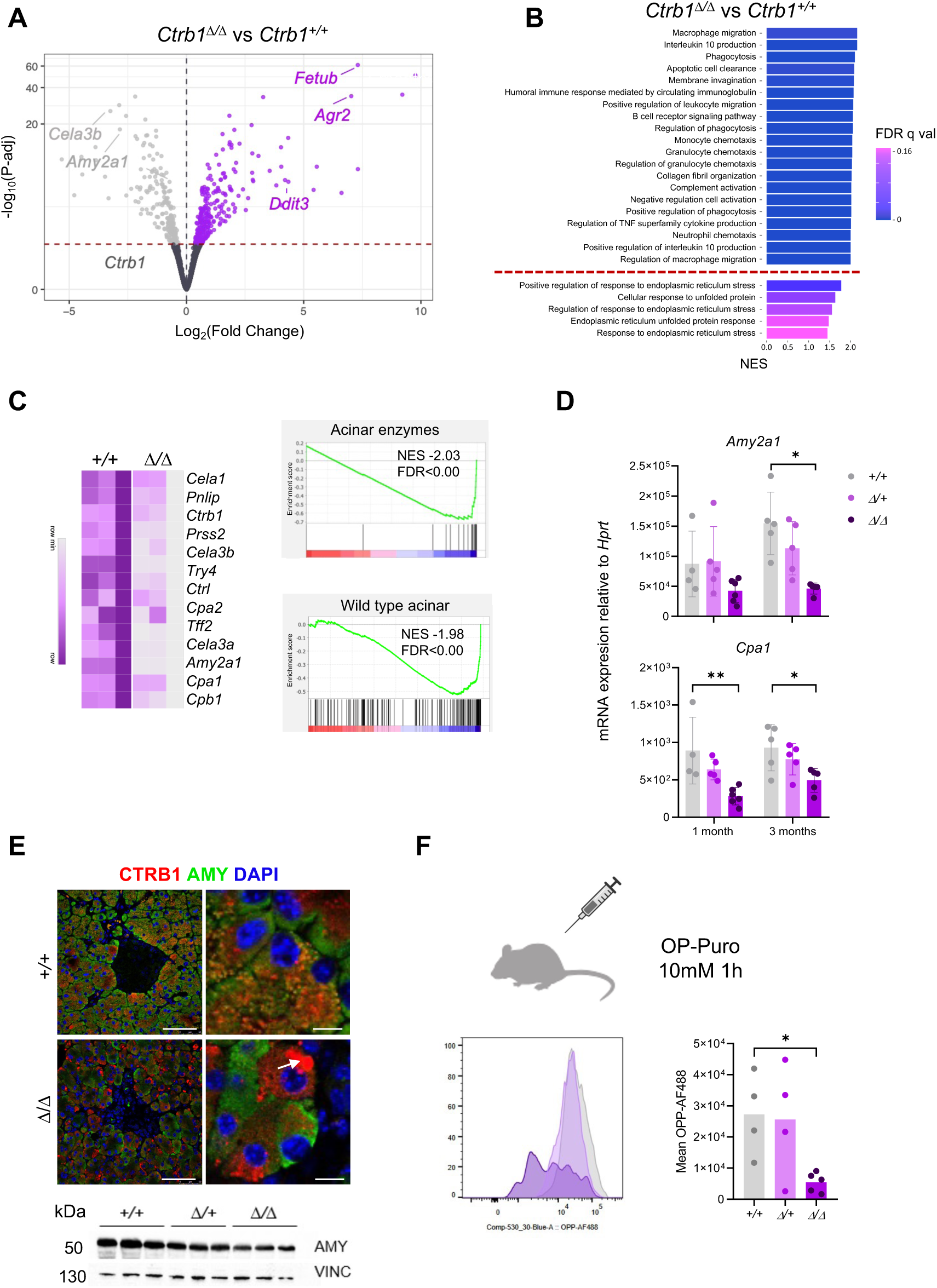
Transcriptomic analysis of pancreata from 3 month-old *Ctrb1^△/△^* and *Ctrb1^+/+^* mice reveals down-regulation of the acinar program and up-regulation of ER stress and inflammatory pathways. **A**. Volcano plot showing differentially expressed genes (P-adj<0.05) in *Ctrb1^△/△^ vs Ctrb1^+/+^* pancreata. Up-regulated genes are labeled in purple and down-regulated genes in light gray. Dashed red line represents the P-adj=0.05 cutoff. Highlighted are exemplary genes related to ER stress and acinar function (n=6 mice/group) **B**. Top 20 significant up-regulated pathways are enriched in inflammatory processes. Red dashed line indicates skipping of pathways to display significant ER stress pathways. **C.** ssGSEA showing a down-regulation of the acinar program at the signature level ^24^ and enrichment plots showing down-regulation of an acinar signature obtained from single cell RNA-Seq data ^25^. **D**. RT-qPCR validation of the down-regulation of *Amy2a1* and *Cpa1* transcripts (n≥4mice/group). **E**. Double IF of CTRB1 and AMY in WT and *Ctrb1^Δ/Δ^* pancreata. In *Ctrb1^Δ/Δ^*mice, CTRB1+ aggregates (white arrow) tend to segregate from AMY-containing structures. Scale bar, 50µm. Western blot shows a decreased expression of AMY in *Ctrb1^Δ/Δ^* pancreata (n=3 mice/group). **F**. Schematic representation of the *in vivo* OP-Puromycin assay to evaluate protein synthesis. Mice were injected with OP-P i.p. and sacrificed 1h later. Reduced protein synthesis in the pancreas of *Ctrb1^Δ/Δ^* mice (n≥4 mice/group).

### Ctrb1^Δ/Δ^ pancreata exhibit down-regulation of the acinar programme and reduced protein synthesis

Because the more common pathway signatures do not contain a specific gene set corresponding to the acinar digestive enzymes and processing machinery, we tested the signature identified by Masui *et al*. as being dependent on the PTF1 component RBPJL ^24^. We noted a highly significant decrease in the activity of this pathway in *Ctrb1^Δ/Δ^* pancreata compared to the WT counterpart (NES = -2.03, FDR q<0.001); similar results were obtained with an acinar signature derived from single cell RNA-Seq (NES = -1.98, FDR q<0.001) ^25^ (**Figure 4C**). These findings were validated by RT-qPCR of *Cpa1* and *Amy2a1* transcripts; reduced expression of AMY in mutant mice was confirmed using IF and western blotting (**Figure 4D, E**). At the protein level, CTRB1 and AMY2A1 co-localized in WT acinar cells but displayed more variable patterns of expression in *Ctrb1^Δ/Δ^* acinar cells (**Figure 4E**).

The 25 top genes expressed in acinar cells account for approximately 80% of all protein-coding transcripts in the pancreas, suggesting that reduced activity of the acinar program would result in major changes in protein synthesis. We administered O-propargyl-puromycin (OP-P) to measure protein synthesis *in vivo*. Similar levels of OP-P uptake were found in WT and *Ctrb1^+/Δ^* mice; in contrast, a significant decrease was observed in *Ctrb1^Δ/Δ^* mice (**Figure 4F**). We also found significantly lower levels of mature 18S rRNA, 5.8S rRNA, and 5.8S splice junction transcripts in the pancreas of 3-month old *Ctrb1^Δ/Δ^* mice (**Supplemental Figure 4**). These results suggest a dramatic dysregulation of key pathways involved in acinar cell identity, ribosome biogenesis, and protein synthesis in homozygous mutant mice.

### Up-regulation of ER stress and UPR-related pathways in Ctrb1^Δ/Δ^ pancreata

The most significantly up-regulated pathways identified were related to ER stress, chaperones, or the unfolded protein response (UPR) (**Figure 5A and Supplemental Table 2**). The three UPR branches are interconnected but a few genes have been proposed to be more closely related to each of them ^26^. **Figures 5A** and **5B** show that the activity of all three branches was significantly up-regulated in *Ctrb1^Δ/Δ^* pancreata both at the gene and pathway levels. Consistently, we noted an overexpression of the UPR markers spliced XBP1 (XBP1-s), BiP, CHOP, and ATF4 using IHC and/or western blotting in *Ctrb1^Δ/Δ^* pancreata (**Figure 5C,D**).

**Figure 5.**
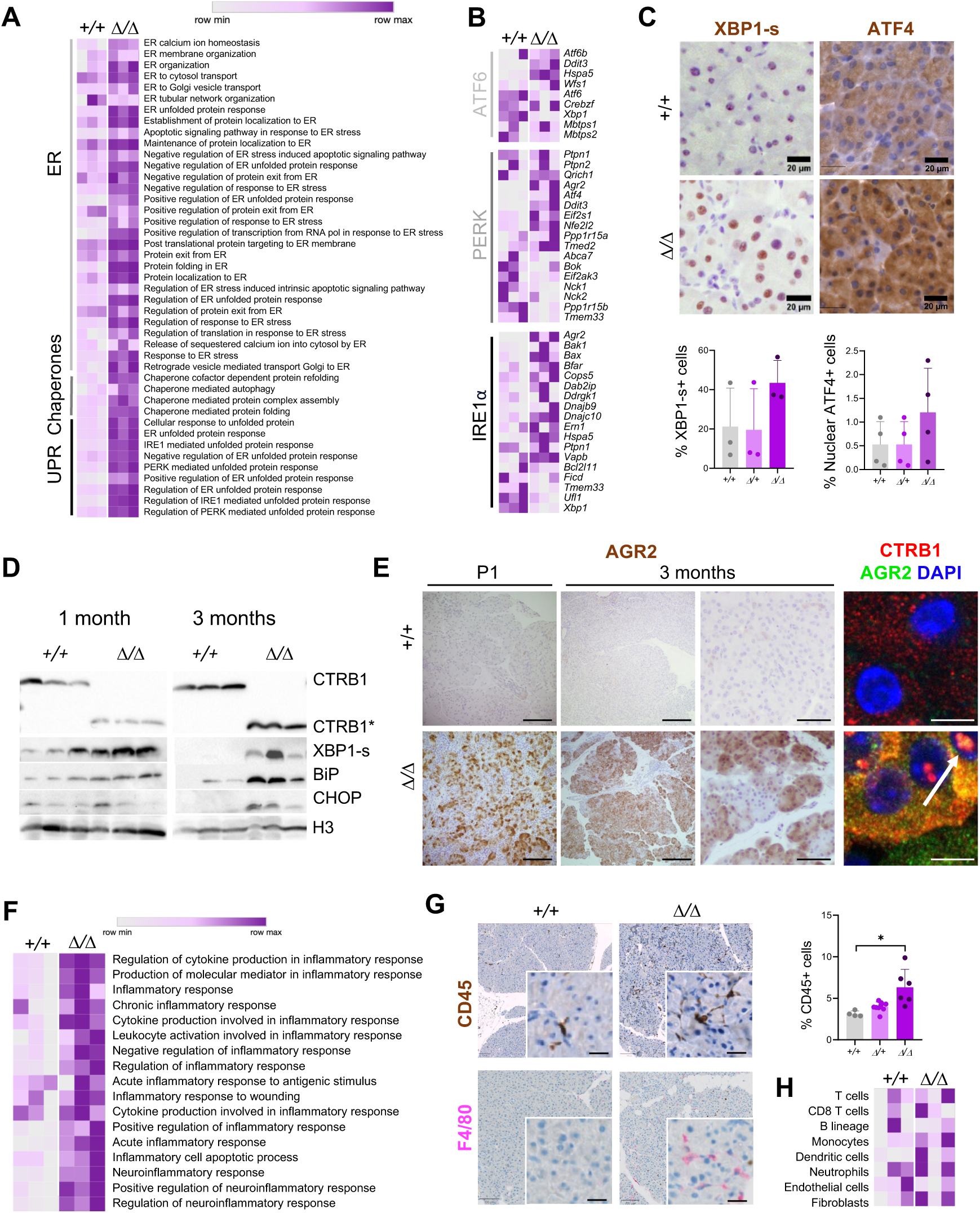
*Ctrb1^△/△^* pancreata show high levels of ER stress and inflammation transcriptomic pathways. **A.** ssGSEA shows that *Ctrb1^△/△^* pancreata display an up-regulation of ER stress-related pathways. **B**. ssGSEA displaying the genes of the three UPR branches. **C**. IHC analysis of ER stress markers (n=3 mice/group). Each dot represents the percentage of positive cells in a full pancreas section. **D.** Western blot showing increased expression of selected UPR markers (n=3 mice/group). **E**. Representative images showing that AGR2 is undetectable in WT pancreas, but it is overexpressed in acinar cells from *Ctrb1^△/△^* mice starting at P1. Double IF of CTRB1 and AGR2 in WT and *Ctrb1^Δ/Δ^* pancreata shows that, in *Ctrb1^Δ/Δ^* mice, CTRB1 and AGR2 are co-expressed in the nucleus (white arrow). Scale bar IHC, 100µm; inset IHC, 50 µm. Scale bar IF, 5µm. **F**. ssGSEA displaying an enrichment of inflammatory pathways. **G**. Significant increase in the number of CD45+ cells in the pancreas of *Ctrb1^Δ/Δ^* mice and representative image showing an increased number of F4/80+ cells (n≥4 mice/group). Each dot represents the percentage of positive cells in a full pancreas section. Scale bar, 100µm; inset, 20µm. **H.** Heatmap showing MCPcounter immune cell deconvolution of RNA-seq data.

One of the most up-regulated genes in mutant mice was *Agr2*, which encodes a protein from the disulfide isomerase family (PDI) that has been linked to cancer and other pathological processes, ^27, 28^. AGR2 is undetectable at the transcript and protein level in normal mouse pancreas. However, its expression was consistently and strongly up-regulated both in *Ctrb1^+/Δ^* and *Ctrb1^Δ/Δ^* pancreata (**Figure 4A** and **Supplemental Figure 3**). IHC revealed a striking mosaic expression pattern in acinar cells with approximately 80% of AGR2+ cells, the remaining cells being unreactive (**Figure 5E**), suggesting a heterogeneous response of acinar cells to the effects of truncated CTRB1. AGR2 also co-localized with CTRB1 in the nucleus of acinar cells (**Figure 5E**).

### Up-regulation of inflammatory pathways in Ctrb1^Δ/Δ^ pancreata

A third group of significantly up-regulated pathways were related to inflammation, including macrophage migration, phagocytosis, and humoral immune response (**Figure 5F and Supplemental Table 2**). IHC analysis confirmed a significantly higher density of CD45+ cells in *Ctrb1^Δ/Δ^* pancreata with higher, non-significant, density in *Ctrb1^+/Δ^* mice (**Figure 5G**). F4/80 expression and MCPCounter deconvolution revealed increased occurrence of monocytes/macrophages and dendritic cells in *Ctrb1^Δ/Δ^* pancreata (**Figure 5G,H**).

### Milder transcriptomic changes in Ctrb1^+/Δ^ pancreata

We also compared the transcriptome of *Ctrb1^+/Δ^*and *Ctrb1^+/+^* pancreata. The number of DEG was lower in the *Ctrb1^+/Δ^* comparison (152 up-regulated and 29 down-regulated) (**Supplemental Figure 3A,B**). Among the most dysregulated genes were several related to ER stress and digestive enzymes (e.g., *Agr2, Hpsa5, Cela3a*) (**Supplemental Table 2**). Several pathways related to protein modification were enriched at nominal p-value<0.05 **(Supplemental Figure 3C**).

### *Ctrb1^Δ/Δ^* pancreata display increased expression of DNA damage response markers

The extensive ultrastructural changes and the up-regulation of pathways associated with DNA damage and apoptosis in *Ctrb1^Δ/Δ^* pancreata (**Figure 6A**) suggest compromised acinar cell survival. Consistently, we found a significantly increased proportion of gH2AX+ acinar cells, as well increased expression of p53, its target p21, and cleaved caspase 3 (**Figure 6B**). This was associated with a significantly higher proportion of Ki67+ acinar cells (**Figure 6C**). These findings support an increased turnover of acinar cells in mutant mice.

**Figure 6.**
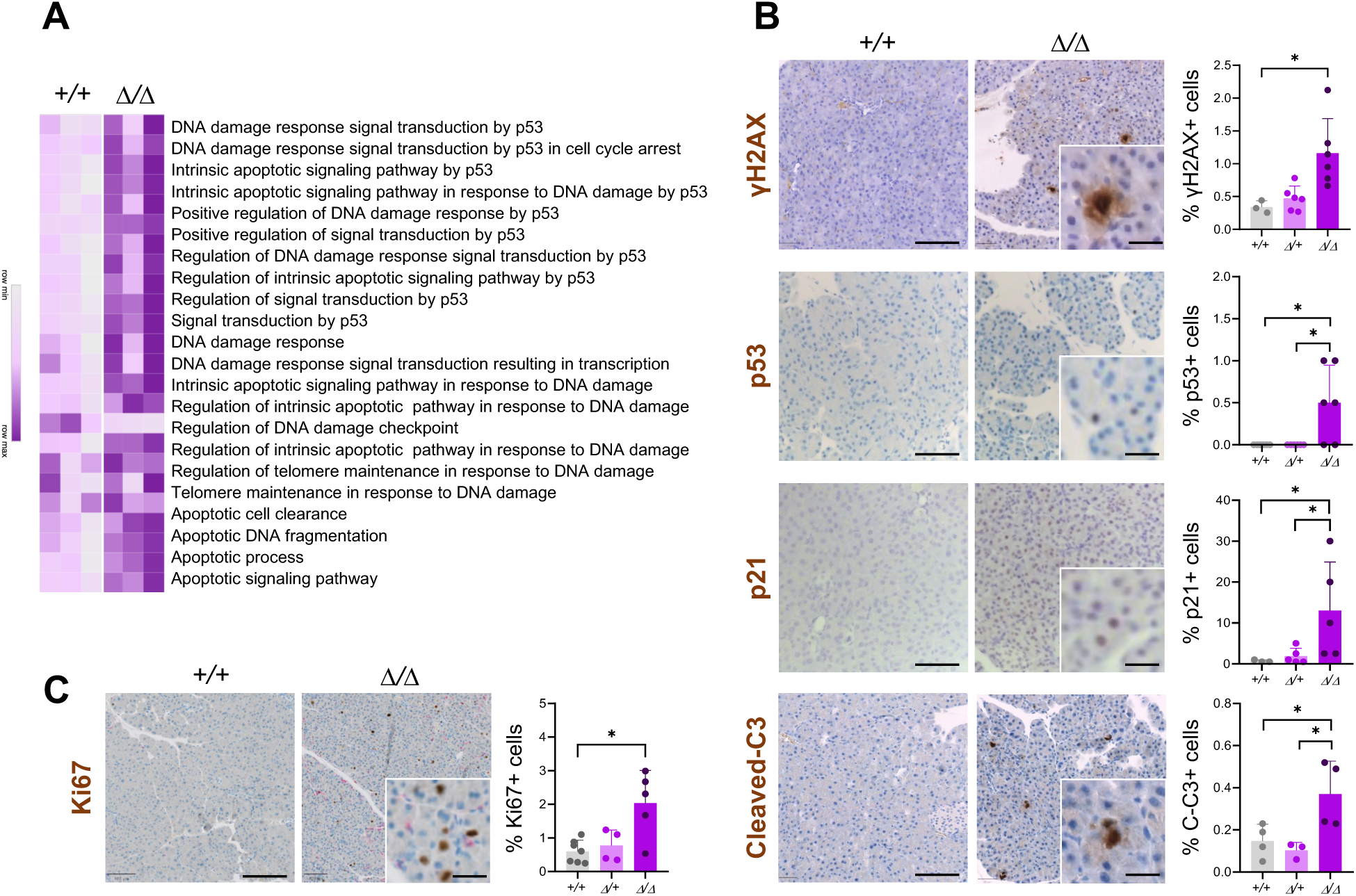
DNA damage and cell cycle markers are up-regulated in the pancreas of *Ctrb1^△/△^* mice. **A.** ssGSEA showing enrichment of DNA damage-related pathways in *Ctrb1^△/△^* mice (n=6 mice/group). **B.** Increased expression of γH2AX, p53, p21, and cleaved caspase 3 in *Ctrb1^△/△^* pancreata. **C**. Increased occurrence of Ki67+ cells in *Ctrb1^Δ/Δ^* pancreata (n≥3 mice/group). Each dot represents the percentage of positive cells in a full pancreas section. Scale bar, 100µm; inset, 20 µm.

### *Ctrb1^Δ/Δ^* mice are more susceptible to damage

To assess whether *Ctrb1^Δ/Δ^* mice are more susceptible to damage, we induced a mild 7-hourly acute caerulein pancreatitis. Mice were sacrificed at 2, 5, and 14 days. Histological examination at day 2 revealed similar histoscores [edema, immune cell infiltration, and acinar-ductal metaplasia (ADM)] in mice of all genotypes (**Figure 7A**). At day 5, partial recovery was observed only in WT mice. WT mice had completely recovered by day 14 whereas *Ctrb1^Δ/Δ^* mice continued to display significantly more damage (**Figure 7A**). IHC analysis confirmed significantly higher CD45+ and F4/80+ cell infiltration (**Figure 7B,C**) and fibrosis, assessed by Syrius red histochemistry (**Figure 7D**) in *Ctrb1^Δ/Δ^* pancreata. Higher Ki67 expression was observed at 48h in *Ctrb1^Δ/Δ^* mice (**Figure 7C**). Both *Ctrb1^Δ/Δ^ and Ctrb1^Δ/+^* showed increased expression of AGR2, KRT19 and SMA after caerulein administration (**Supplemental Figure 5A-C).** As in basal conditions, the phenotype of *Ctrb1^Δ/+^* mice was milder.

**Figure 7.**
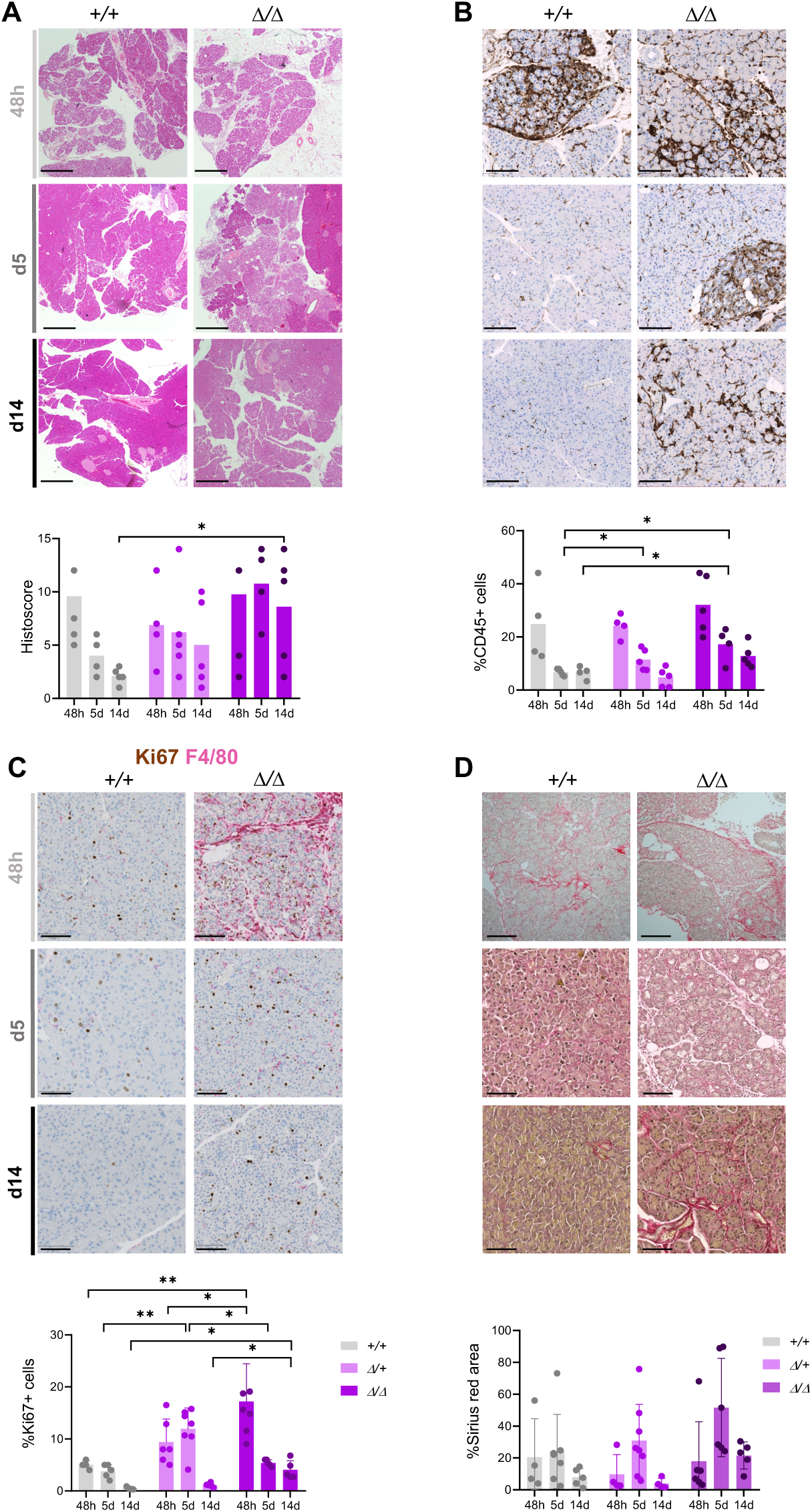
*Ctrb1^△/△^* mice show impaired recovery upon induction of an acute caerulein pancreatitis. **A.** Histological analysis shows similar levels of damage in WT and mutant mice at 48h, with impaired recovery in *Ctrb1^△/△^* mice at days 5 and 14 (n≥4 mice/group). Scale bar, 500µm. **B**. Increased number of CD45+ cells in *Ctrb1^△/△^* pancreata at 5 and 14 days (n≥4 mice/group). Each dot represents the percentage of positive cells in a full pancreas section. Scale bar, 100µm. **C.** Increased F4/80 infiltration in *Ctrb1^△/△^* pancreata at 48h and 5 days. Increased Ki67 expression in *Ctrb1^△/+^* mice at day 5 and, in *Ctrb1^△/△^* mice, at 48h and 14 days (n≥4 mice/group). Scale bar, 100µm. **D**. Sirius Red staining and quantification reveals a peak in fibrosis at day 5 in *Ctrb1^△/△^* mice that is not resolved by day 14 (n≥4 mice/group). Scale bar, 100µm.

### TUDCA and Sulindac alleviate the ER stress caused by mutant CTRB1

Several FDA-approved drugs are available with proven ability to mitigate ER stress, including TUDCA, a chaperone-like molecule ^29^. We administered TUDCA i.p. to 3 month-old mutant mice for 4 weeks (**Figure 8A**). The pancreas of TUDCA-treated mice showed normal histology (**Figure 8B**). Ultrastructural analysis revealed a reduction in ER stress indicators, including more abundant ER stacks of normal, flattened, appearance (**Figure 8C**). Expression of UPR-related genes in TUDCA-treated *Ctrb1^Δ/Δ^* mice (e.g., *BiP, Chop, Agr2, Xbp1,* and *Xbp1-s*) was reduced at borderline significance (for *BiP, Agr2, Xbp1,* and *Xbp1-s*, p=0.057; for *Chop,* p=0.114) (**Figure 8D**). A similarly reduced expression of BiP, CHOP, and ATF4 was observed using IF (**Figure 8G**), suggesting a partial improvement of the phenotype.

**Figure 8.**
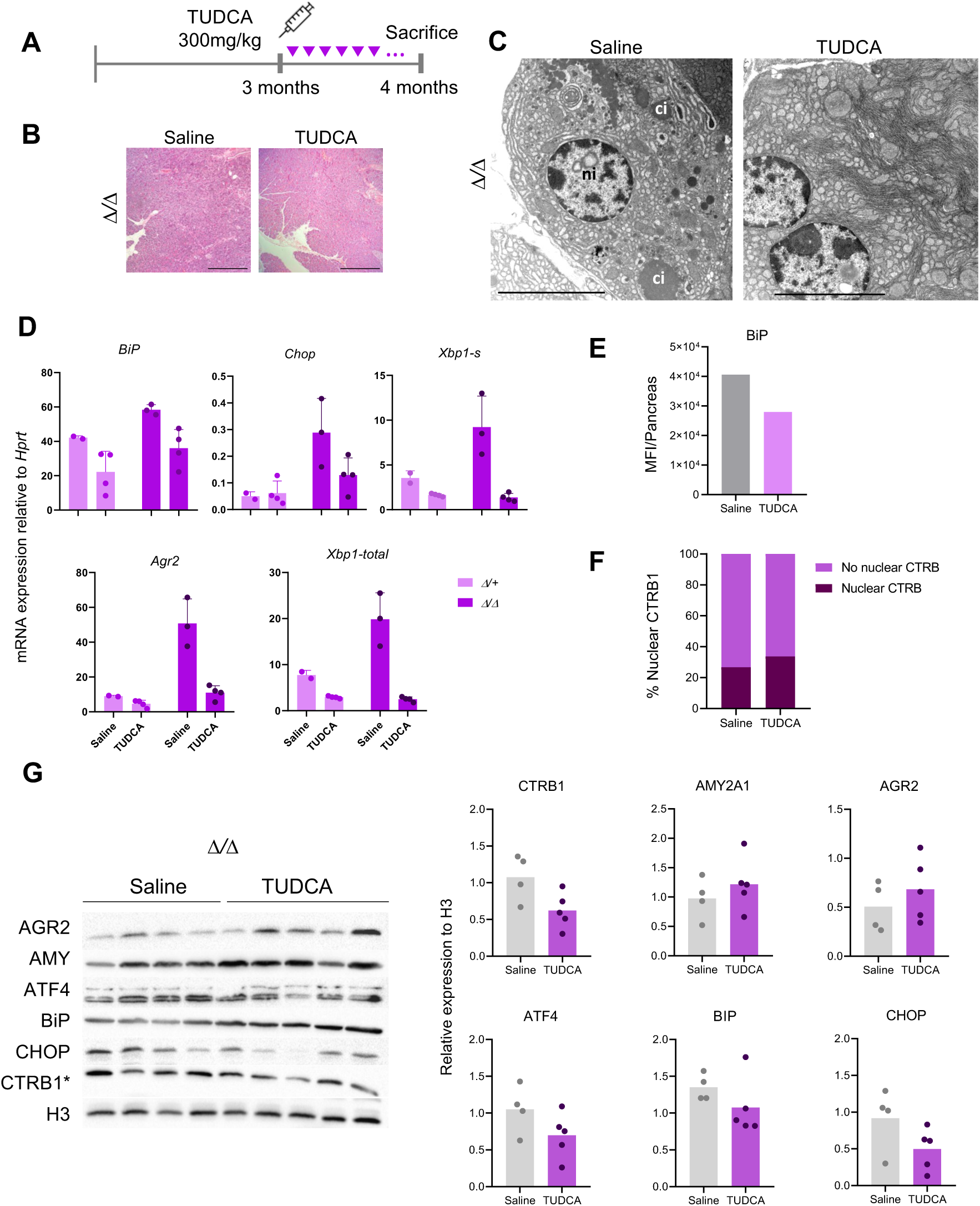
TUDCA administration to *Ctrb1^Δ/Δ^* mice results in a mild amelioration of ER stress features. **A.** Schematic representation of TUDCA administration. **B**. Histological analysis reveals normal histology in control and TUDCA-treated *Ctrb1^Δ/Δ^* pancreata (n≥4 mice/group) Scale bar, 500µm. **C**. Ultrastructural analysis reveals more prominent, well-organized, ER cisternal stacks in acinar cells of TUDCA-treated mice. **D**. RT-qPCR analysis reveals reduced expression of ER stress markers in TUDCA-treated mice (n≥2 mice/group). **E**. IF analysis reveals a slight decrease in BiP expression in the pancreas of TUDCA-treated mice (n=2 mice/group). **F**. Nuclear CTRB1 accumulation in acinar cells of TUDCA-treated mice (n=2 mice/group). **G.** Western blot analysis of CTRB1, amylase, and ER stress-related proteins reveals reduced expression of CTRB1, ATF4, CHOP, and BIP in TUDCA-treated mice (n≥4 mice/group).

Given the up-regulation of inflammatory pathways, we also tested the effect of Sulindac, a non-selective COX inhibitor. Six week-old mice received Sulindac for 6 weeks (**Supplemental Figure 6A**), after which the pancreas displayed a normal histology (**Supplemental Figure 6B**). A decrease in the expression of ER stress markers (*BiP*, p=0.015; *Chop*, p=0.008), but not of *Amy2a1* or *Cd45,* was observed (**Supplemental Figure 6C**). At the protein level, there was a mild reduction in AGR2 expression in *Ctrb1^Δ/+^* mice receiving Sulindac. Ki67 and CD45 expression were similar in both conditions (**Supplemental Figure 6D,E**).

### *Ctrb1^Δ/Δ^* mice recapitulate transcriptomic features of the pancreas of human carriers of the *CTRB2* exon 6 deletion

To determine whether our mouse model recapitulates features of human carriers of the *CTRB2* exon 6 deletion, we mined publicly available data from the pancreas of healthy individuals from the Genotype-Tissue Expression (GTEx v8) project. After rigorous imputation of the *CTRB2* exon 6 deletion and transcriptomic analysis of exon-exon junction counts (for details, see **Supplemental Material**), we focused on 159 individuals. These included 17 *CTRB2^Δ/+^* and 2 *CTRB2^Δ/Δ^* subjects (**Figure 9A**). A signature extracted from the mouse RNA-seq data comprising the 291 significantly up-regulated genes in *Ctrb1^Δ/Δ^* pancreata was significantly up-regulated/enriched in the samples from exon 6 deletion homozygous allele carrier subjects (**Figure 9B**). Differential expression analysis comparing homozygous to wild type individuals revealed an enrichment of ER stress-related pathways, as previously reported ^21^ (**Figure 9C**). These findings underscore the relevance of our mouse model to human disease.

**Figure 9.**
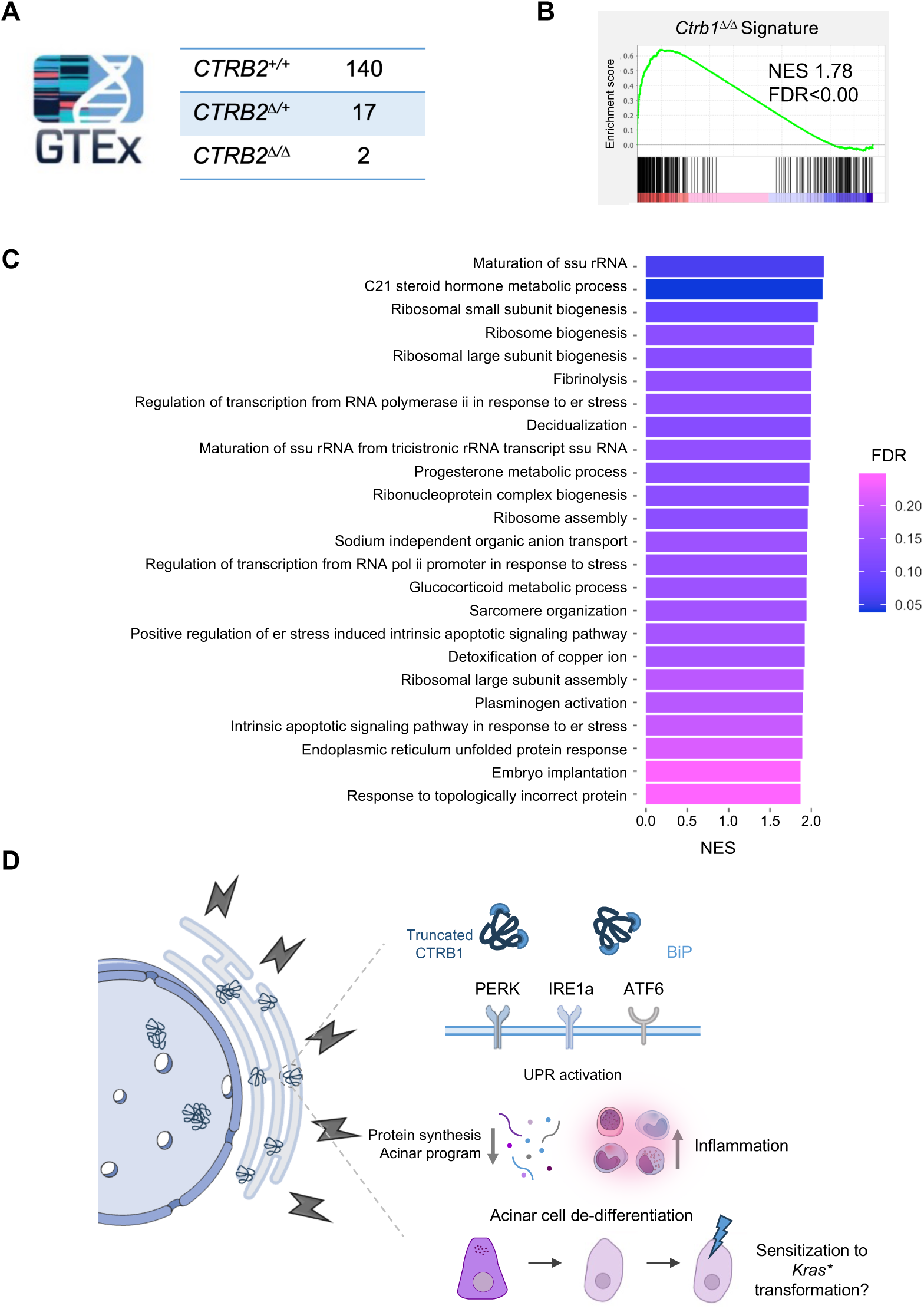
Enrichment of the *Ctrb1^△/△^* transcriptomic signature in the normal pancreas of human GTEx *CTRB2^△/△^* samples. **A**. Summary of the genotype distribution resulting from the overlap between the published data ^21^ and our imputation of the exon 6 deletion in subjects from the GTEx cohort. **B.** GSEA plot showing that homozygous samples are enriched in the *△/△* signature (NES=1.8, FDR < 0.00). The signature includes the 291 significantly up-regulated genes identified in the *Ctrb1^△/△^ vs Ctrb1^+/+^* RNA-seq comparison. **C**. Pathways enriched in samples from *CTRB2^△/△^* and *vs CTRB2^+/+^* subjects include those related to ER stress. **D**. Proposal of a model for the effects of the exon 6 deletion based on the work reported in mice. Synthesis of mutant CTRB1 results in an overload of misfolded proteins that results in activation of ER stress and UPR pathways. This leads, through yet undefined mechanisms, to a down-regulation of the acinar program associated with reduced total protein synthesis and an up-regulation of inflammatory pathways. The observed effects are most severe in *Ctrb1^△/△^* mice; an intermediate phenotype is observed in *Ctrb1^△/+^* mice. The attenuation of the acinar cell differentiation program may sensitize these cells to mutant *Kras*-driven transformation.

## DISCUSSION

GWAS provides unbiased approach to identify associations of genotypes with phenotypes; however, by themselves they cannot provide direct causal evidence explaining the underlying molecular mechanism of risk. Here, we build on the fine mapping of a GWAS signal at chr16q23.1 pointing to a deletion overlapping exon 6 of *CTRB2* as the variant underlying the PDAC risk signal. We report a new mouse model mimicking this variant and providing strong evidence supporting its causal involvement in pancreatic pathophysiology.

Using CRISPR-Cas9, we targeted the human exon 6 deletion to the mouse orthologue *Ctrb1* and show that it leads to the synthesis of a truncated CTRB1 protein that is mainly present in the insoluble fraction of pancreatic lysates and forms aggregates in acinar cells. Several important results from our study provide evidence about how this variant may contribute to disease.

Despite massive disruption of acinar cell homeostasis demonstrated by ultrastructural analysis, standard histology was inconspicuous and we did not observe ADM, a characteristic response of acinar cells upon stress. In agreement with the normal histology, homozygous mutant mice are viable, breed normally, and show no signs of pancreatic dysfunction including exocrine failure. As predicted, we observed reduced chymotrypsin activity in homozygous mutant mice, indicating that the phenotype does not result from a gain-of-function. The ultrastructural analysis revealed a dramatic dilation of ER cisternae, including the nuclear envelope, and intracisternal granules (ICG). An increase in ICG has been reported upon starvation or cobalt addition to acinar cells, associated with an increased concentration of digestive enzymes in the rough ER. These ICGs are massive aggregates of misfolded proteins that exclude BiP and PDI ^30^. Accordingly, RNA-Seq analysis disclosed an up-regulated expression of multiple genes (e.g., *Ddit3/Chop*, *Agr2*, *BiP*) and pathways related to the UPR and ER stress response. We found evidence for involvement of the three branches of the UPR; broad UPR activation is likely responsible for the down-regulation of protein synthesis observed *in vivo* in *Ctrb1^Δ/Δ^*pancreata. *Agr2* mRNA was undetectable in WT pancreata in basal conditions and *Agr2* was one of the most significantly up-regulated genes both in heterozygous and homozygous mutant mice. In contrast, *Agr2* transcripts are undetectable upon induction of pancreatitis with caerulein or reprogramming with OSKM ^25^ and AGR2 expression is not associated with ADM, as shown in **Supplemental Figure 5**. AGR2 is linked to the activation of the IRE1 and ATF6 arms of the UPR ^28, 31^ and its participation in PDAC initiation and progression has been proposed ^32^. *AGR2*-silenced cells display reduced ER quality control and folding capacity and are highly sensitive to ER stress ^32^. AGR2 selectively represses the endonuclease activity of IRE1b - but not IRE1a - in colonic goblet cells ^33, 34^. However, single cell RNA-Seq shows that IRE1b is undetectable in acinar cells, pointing to a conserved role as part of the UPR in but a different mechanism of action in colonocytes and acinar cells.

AGR2 immunostaining showed a striking mosaic pattern, suggesting a heterogeneous response of acinar cells to misfolded CTRB1. We propose three scenarios to explain this finding. First, it may result from differential activation of the three UPR branches in acinar cells. Second, it may result from dynamic regulation of the response of cells to misfolded protein accumulation. Mutant CTRB1 accumulation, activation of the ER stress and UPR to restore homeostasis, partial ER stress resolution, and deactivation of the UPR, followed by an up-regulation of the acinar program might contribute to pathological homeostasis. This on/off response may occur at different rates in different cells, resulting in a mosaic-like phenotype. Finally, the mosaic pattern may reflect acinar cell heterogeneity in the adult pancreas ^35, 36^.

While a few other genetic mouse models of misfolded protein-driven pancreatic ER stress have been reported, they implicate different molecular mechanisms. One group refers to gain-of-function mutations in pancreatic genes such as *Cpa1*^19^ or *Pnlip*^37^. These strains mimic rare mutations in these genes in patients with hereditary pancreatitis. Another group involves strains with mutations in genes that are involved either in general ER function or in secretion and are widely expressed in normal tissues (e.g., *Perk*, *Mist*, *Xbp1*, *Sec3b*) ^38–42^, some of them being neonatal lethal (e.g., *Sec3b*). A third group relates to strains with pancreas-specific deletion of autophagy genes: *Atg5* or *Atg7* knockout mice share some features (e.g., reduced protein synthesis) with our model but they have a more severe basal phenotype ^43,44^. *Ccpg1* hypomorphic mice have an ER-selective autophagy defect and a milder phenotype ^45^. We did not observe ultrastructural or transcriptomic evidence of the activation of autophagy in *Ctrb1^Δ/Δ^* mice, an intriguing finding considering the massive ER dilation. However, there was a slight increase in p62 expression in *Ctrb1^Δ/Δ^*pancreata (not shown). These findings support the unique phenotype of our model.

An intriguing ultrastructural observation was the presence of large nuclear inclusions in a fraction of acinar cells, sometimes in association with signs of cellular stress and/or apoptosis. IF analysis strongly suggested that they correspond to protein aggregates containing misfolded CTRB1 and chaperones, such as AGR2 and BiP. It is tempting to speculate that the inclusions result from the rupture of the nuclear envelope - as also observed in the cytosol in the proximity of ruptured ER membranes. Whether these aggregates impact on genome organization and/or function requires further study. Nuclear protein aggregates have been shown to disrupt the distribution of nuclear compartments, altering chromatin accessibility, transcription, and RNA processing ^46^.

As in other genetic mouse models of pancreatic disease, the mild basal phenotype is accompanied by a highly sensitized state to damage (e.g., acute pancreatitis). Mice in which *Gata6* is inactivated in the embryonic pancreas display a largely normal histology at 10-12 weeks-age but they do not recover properly from a mild acute caerulein-induced pancreatitis ^12^. A similar phenotype is observed in *Nfic*-null mice and in other models ^14^. In these cases, damage scores are similar in WT and mutant mice, but a defect in recovery is apparent. In other instances, such as *Nr5a2* heterozygous mice, both increased damage and delayed recovery occur ^11,13^.

*Ctrb1^+/Δ^* mice display an intermediate phenotype between WT and *Ctrb1^Δ/Δ^* mice. In some respects, heterozygous mice more closely resemble WT mice (e.g., chymotrypsin activity or protein synthesis) despite the fact that approximately half of *Ctrb1* transcript expressed lacks exon 6. This may reflect the ability of acinar cells to cope with a considerable increase in misfolded proteins, sufficient to mitigate the phenotype of *Ctrb1^+/Δ^* mice but insufficient to cope with the protein overload present in *Ctrb1^Δ/Δ^* mice. These findings suggest non-linear relationships between the amount of misfolded proteins, the cellular responses (e.g., UPR activation, DNA damage, apoptosis, proliferation), and the biological outputs. Understanding these processes will be key to interpreting quantitative aspects of the GWAS risk associations: in an additive model the odds for PDAC among individuals with one deleted allele is 1.36 whereas in a recessive model the OR is 1.91 ^21^.

Our findings need to be placed in the context of the phenotype of mice in which *Ctrb1* was inactivated. *Ctrb1* knockout mice have reduced chymotrypsin activity and lack pancreatic alterations up to one year of age, but they develop a more severe acute caerulein-induced pancreatitis ^22^, a distinct finding compared to our model where severity was similar, but recovery was impaired.

The mouse strain described here provides the opportunity to assess therapeutic strategies that could be applied in carriers of the *CTRB2* exon 6 deletion variant. To explore these options, we have used TUDCA - an FDA-approved drug for the treatment of primary biliary cholangitis relieving ER stress - and Sulindac, a non-selective COX inhibitor. We noted a modest amelioration of ER stress markers, indicating the potential to impact on the mechanisms involved in the disease. The lesser effects observed in homozygous mutant mice suggests that very high ER stress levels may not be as readily reversible/preventable. The identification of the UPR arm playing a more fundamental role in the phenotype may guide therapeutic optimization.

The ensemble of data reported here support a model whereby accumulation of misfolded CTRB1 in the ER results in activation of multiple UPR pathways and down-regulation of protein synthesis (**Figure 9D**). This was accompanied by a down-regulation of the acinar differentiation programme that may alleviate the protein synthesis load in acinar cells. The resulting ER stress, and the incomplete differentiation of acinar cells, may also lead to the up-regulation of inflammatory pathways ^11–14^. Both the incomplete acinar differentiation and the accompanying inflammation may sensitize the pancreas to signaling by mutant *Kras*, possibly explaining the GWAS association with increased PDAC risk observed in patients.

The enrichment of a *Ctrb1^Δ/Δ^*signature and of ER stress-related signatures in GTEx *CTRB2* exon 6 deletion carriers is a strong indication that this variant is well recapitulated in our model. Our findings strongly support the notion that the *CTRB2* exon 6 deletion is a causal variant for human PDAC. Genetic variation in *CTRB1/CTRB2* has also been associated with an increased risk of chronic pancreatitis, but it is currently unclear whether the *CTRB2* exon 6 deletion is also causal for this association ^20^. The mouse strain reported here provides unique opportunities to understand the interplay between pathological ER stress, inflammation, and cancer and to identify PDAC preventative strategies.

## Acknowledgements

We thank the members of the Epithelial Carcinogenesis Group for discussion and criticism, S. Rueda, E. Andrada, M. Andueza, N. Dusetti, J. Iovanna, S. Lowe and J. Ferrer and their teams for valuable contributions, and the Histopathology Unit of CNIO for technical support.

## Data access

RNA-Seq data have been deposited in GEO with accession number GSE272278.

## Author contributions

Conceptualization: CB, IF, ELM, LTA, NM, and FXR; data acquisition: CB, IF, BC, ML, ELM, NP, and MM; background data acquisition: KEC, JWH, LTA, ELM, NM, and FXR; resource generation: PV and SO; data analysis: CB, IF, BC, ML, JMV, ELM, AC, IP, and RM; data curation: JMV and ELM; writing: CB, IF, ML, ELM, and FXR; supervision: NM and FXR; funding acquisition: RM, NM, and FXR. All authors reviewed the manuscript and provided comments.

