## Supplementary Material for "A common CTRB misfolding variant associated with pancreatic cancer risk causes ER stress and inflammation in mice"

### SUPPLEMENTAL INFORMATION

#### Index

**SUPPLEMENTAL MATERIALS AND METHODS**

Supplemental References

**SUPPLEMENTAL TABLES**

Supplemental Table 1. Top 20 expressed genes in normal mouse and human pancreas

Supplemental Table 2. RNA-Seq analysis: differentially expressed genes (provided as a supplemental Excel file)

Supplemental Table 3. List of antibodies used in this study

Supplemental Table 4. List of oligonucleotide primers used for RT-qPCR in this study

**SUPPLEMENTAL FIGURES**

### SUPPLEMENTAL MATERIALS AND METHODS

**Mouse strains.** *Ctrb1*$^{\Delta ex6}$ mice were maintained in a C57BL/6 background. Experiments were performed using 4–12 week-old mice of both sexes, as indicated in the text. Breedings were set in heterozygosity and littermates were used as controls. Mice of all genotypes were co-housed. All animal procedures were approved by local and regional ethics committees [(Institutional Animal Care and Use Committee and Ethics Committee for Research and Animal Welfare, Instituto de Salud Carlos III) (CBA 09_2015_v2) and Comunidad Autónoma de Madrid (ES280790000186)] and performed according to the European Union guidelines.

A mild acute pancreatitis was induced by 7 hourly injections of the cholecystokinin analog caerulein (Bachem) at 50 µg/kg. In brief, animals were weighed before the procedure and caerulein was administered intraperitoneally. Mice were killed by cervical dislocation after 48 h, 5 and 14 days after the first caerulein injection. Tauroursodeoxicolic acid (TUDCA) (CAS 14605-22-2 – Calbiochem, 580549-5GM) was administered i.p. daily at a dose of 300mg/kg for 30 days to 12 week-old mice. Control mice received PBS. Sulindac (S8139-25G) was administered *ad libitum* in drinking water at a concentration of 200mg/L for 3 months to 6-week-old mice. For most experiments, ≥5 mice per group were used. No specific randomization method was used.

**Tissue collection and preparation.** Mice were sacrificed by cervical dislocation, unless otherwise indicated. A small piece from the tail of the pancreas was flash-frozen and stored at -80 ºC for RNA and protein extraction; the remaining pancreas was fixed overnight in 4% PBS-buffered formaldehyde. After fixation, tissues were embedded in paraffin, serially sectioned (3 µm), and either stained with hematoxylin and eosin (H&E) or used for immunohistochemistry.

**Histological scoring.** H&E-stained sections were used to score edema, leukocyte infiltration and ADM. Scores considered the grade of severity (0-3) and the extent of the feature (0, absent; 1, focal; 2, multifocal; and 3, diffuse). The individual scores were then added into a global histoscore that ranges from 0-18. Lipomatosis was scored based on severity and extent, as described above (0-6). All sections were reviewed blindly by CB, IF, and FXR.

**Immunohistochemical analysis and quantification.** Sections of formalin-fixed paraffin-embedded (FFPE) tissue blocks were heated for 30 min at 60 ºC, deparaffinized with xylol, and rehydrated with an alcohol series and distilled water. Sections were boiled in 10mM citrate buffer (pH=6.0) for 10 min for antigen retrieval. Next, sections were incubated with 3% $H_2O_2$ in methanol for 30 min, washed with PBS, and incubated with 3% BSA in PBS for 1h at room temperature (RT). Primary antibodies diluted in 3% BSA in PBS were added overnight at 4 ºC. Sections were washed three times with PBS/0.1% Triton X-100 and incubated with the corresponding secondary EnVision+ HRP secondary antibodies (Dako). DAB was used as chromogen; nuclei were counterstained for 45 sec with Carazzi's hematoxylin. Primary antibodies and dilutions used are listed in Supplementary Table 3. Images were digitalized using a Mirax 3D Histech scanner and quantified using QuPath 4.3.0.

**Immunofluorescence (IF) analysis and quantification.** For IF staining, sections were handled as described above and were incubated with primary antibodies overnight at 4 °C. For double IF, the corresponding antibodies were added simultaneously and incubated overnight at 4 °C, ensuring that there were no cross-species signals. Sections were then washed with 0.1% Triton–PBS, incubated with the appropriate fluorochrome-conjugated secondary antibody, and nuclei were counterstained with DAPI. After washing with PBS, sections were mounted with Prolong Gold Antifade Reagent (Life Technology). Primary antibodies and dilutions used are listed in Supplementary Table 3. The images were acquired using a TCS SP5 confocal microscope (Leica-Microsystems) with a 63x objective and 488 and 555 laser lines. Image processing was performed using LAS AF and FIJI software.

We employed the Cellpose[1] deep learning framework to train 2D segmentation models for DAPI-stained nuclei, CTRB1 aggregates, and calreticulin-positive structures. Microscopy images were manually annotated to create ground truth masks for each component. Then, the models were fine-tuned using a combination of U-Net architecture and transfer learning, enabling precise quantification and localization of cellular components, which we used for further statistical analyses into cellular dynamics. To assess differences between conditions and time-lapses, we employed a two-sample t-test. Specifically, we calculated p-values to compare number of CTRB1 aggregates ((N), Mean Intensity, Sum Intensity) per cyto-nucleus within the homozygous condition across different time-lapses (1-3). Additionally, we determined p-values to compare CTRB1 aggregates (N, mean intensity, sum intensity) per cyto-nucleus between the homozygous and wild type conditions for all time-lapses combined. Furthermore, within each month (1 and 3), we computed p-values to compare CTRB1 aggregates (N, mean intensity, sum intensity) per cyto-nucleus between the homozygous and wild type conditions.

**Ultrastructural analyses.** For transmission electron microscopy analysis of the pancreas, mice were sacrificed in a $CO_2$ chamber and perfused with 1% glutaraldehyde and 1% paraformaldehyde in 0.12 M phosphate buffer (pH 7.2), using a MINIPULS3 peristaltic pump (Gilson, USA). Whole pancreata were removed, sectioned into small fragments (1mm), and postfixed in the same fixative for 2h at RT. Tissue samples were then washed in 0.12M phosphate buffer (3x15 min) and postfixed in 2% osmium tetroxide for 2-4h at RT. Tissue fragments were dehydrated in an acetone series, embedded in araldite resin (DURCUPAN, Sigma-Aldrich, USA) and baked at 60ºC for 3 days. Hardened blocks were cut with a 35° diamond knife (Diatome, USA) using an UltraCut UC7 ultramicrotome (Leica Microsystems, Germany). Semithin sections (1µm thick) stained with toluidine blue were used for light microscopy tissue analysis. Ultrathin sections (50-70 nm) were collected on formvar-coated copper grids and contrast-stained with uranyl acetate and lead citrate. They were examined with an JEM 1011 (JEOL, Japan) electron microscope, operating at 80kV. Micrographs were taken with a camera (Orius 1200A; Gatan, USA) using the DigitalMicrograph software package (Gatan, USA). Electron micrographs were processed using Adobe Photoshop CS6 (13.0.1) (Adobe Systems).

**Protein extraction and western blotting.** Tissue was lysed on ice-cold RIPA buffer [pH 7.4, containing 50 mM Tris-HCl, pH 8.0, 150 mM NaCl, 1% NP-40, 0.5% sodium deoxycholate, 5 mM EDTA, 0.1% SDS, 2 mM $Na_3VO_4$, 10 mM NaF, 10 mM $Na_4P_2O_7$, protease inhibitor cocktail (Sigma, catalogue no., P8340), phosphatase inhibitor 2 (Sigma, catalogue no. P5726) PVDF and sodium ortovanadate] for 30 min at 4 °C, and sonicated with a Branson Sonifier 250. Lysates were centrifuged at 6,000 rpm for 5 min; protein concentration of the supernatant was quantified using the DC Protein Assay Reagent kit (Bio-Rad).

Proteins (30-50 µg) were resolved by 12-15% SDS-PAGE and transferred to nitrocellulose membranes (Amersham). After blocking in 1X PBS containing 5% (w/v) skim milk and 0.1% Tween20 (v/v), membranes were incubated with the primary antibody diluted in binding buffer [1X TBS with 5% (w/v) skin milk and 0.1% Tween20 (v/v)] overnight at 4 °C. Membranes were washed 3 times with 1X TBS containing 0.1% Tween20 (v/v), incubated with horseradish peroxidase–conjugated rabbit anti-goat or goat anti-mouse Ig (Sigma-Aldrich), and washed toReactions were developed with Immobilon® crescendo Western HRP substrate (Millipore) and imaged in a ChemiDoc Imaging system (Bio-Rad). Primary antibodies and dilutions used are listed in Supplementary Table 3. Densitometry analysis of digitalized western blotting images was performed using Fiji software.

To assess the solubility of proteins, the detergent-insoluble fraction was washed twice with cold PBS, resuspended in Laemmli buffer, and heated at 96 °C for 5 min. The supernatant was mixed with 1× Laemmli buffer and heated at 56 °C for 15 min after total protein concentration had been determined using the Pierce BCA Protein Assay Kit (Thermo Fisher

Scientific, catalogue no. 23225). Soluble and insoluble fractions were resolved by 12% SDS-PAGE as previously described.

**Protein aggregation assay.** Protein aggregates were measured using the Proteostat protein aggregation assay (ENZ-51023). Shorty, 2 µL of the diluted Proteostat ® Detection Reagent was added into the bottom of each well of a 96-well microplate. Then the protein mix of interest (98 µL) was added to each well at a concentration of 10 µg/mL. The microplate containing test samples was incubated in the dark for 15 min at RT and the signal generated was read with a fluorescence microplate reader VICTOR Nivo ™ Multimode Plate Reader, PerkinElmer using an excitation setting of 550 nm and an emission filter of 600 nm.

**Protein synthesis analysis.** Mice were injected with OP-P (49.5 mg OP-Puro per kilogram of body weight). After 1 h, mice where sacrificed and the pancreas was collected and minced in cold HBSS. Pancreas was then digested [1 mg/mL collagenase V (Sigma) + 0.1 mg/mL DNase I + 2 U/mL Dispase II (Roche) + 0.1 mg/mL Soybean Trypsin Inhibitor (Life Technologies - 17075- 029) in HBSS (with Ca2+/Mg2+; Invitrogen, # 24020)], filtered through 0.22 PES Steriflip (Millipore), and incubated for 42 min at 37 ºC in a GentleMacs dissociator. Afterwards, pancreata were washed with PBS and 0.05% trypsin-EDTA was added for 3 min at 37 ºC. Then, pancreata were washed with FACS buffer (10mM EGTA, 2% FBS in CA/Mg-free PBS), centrifuged, and resuspended in FACS buffer. Cells were passed through a 100 µm strainer, centrifuged, and stained with LIVE/DEAD® FIXABLE AQUA DEAD CELL STAIN (ref. L34965, Life technologies) and CD45 following manufacturer's instructions. Then, cells were fixed and permeabilized, resuspended in 3.7% PFA, and incubated on ice protected from light for 15 min. After washing with cold PBS 1% BSA and centrifugation, cells were permeabilized with 0.5% Triton X-100 in PBS and incubated for 15 min at RT. Then, CLICK-IT PLUS OPP AF488 was used following manufacturer's instructions. Data was acquired by flow cytometry in a FACS Canto II cytometer.

**Serum amylase quantification.** Serum samples were diluted 1:50 in distilled water. Amylase quantification was performed in an ABX Pentra automated clinical chemistry analyser (HORIBA ABX) using Amylase CP diagnostic reagents (A11A01628, HORIBA ABX).

**RNA isolation from total pancreas and real-time quantitative PCR.** RNA was isolated from pancreas as described elsewhere [2]. Briefly, total RNA was isolated using the phenol-chloroform method. RNA was treated with DNase I (DNA-free DNase Treatment & Removal Reagents, Ambion) according to manufacturer's instructions. For gene expression, cDNA was synthesized using reverse transcriptase and random hexamer primers, according to the manufacturer's protocols. RT-qPCR assays were performed on a StepOne Real-Time PCR System using Standard SYBR Green Master Mix. The primers used for RT-qPCR assays are specified Supplementary Table 4.

**RNA-sequencing (RNA-seq) analysis of mouse pancreata.** RNA quality was determined with the RNA Integrity Number (RIN) using an Agilent 2100 Bioanalyazer. RIN- and gender-matched samples with RIN values ranging from 5.7 to 8.3 were sent to the CNIO Genomics Unit for library preparation and sequencing (N=6 mice/genotype sequenced as pools of two mice of the same genotype). RNA-seq data was processed using **cluster_rnaseq** (https://github.com/cnio-bu/cluster_rnaseq) using default parameters. This software enables evaluating read quality, aligning reads to the mouse reference genome (mm10), and performing differential expression analysis using DEseq to obtain a list of deregulated genes between samples. Statistical significance for differential gene expresion (DEG) was set at p-adj < 0.05. The matrix of normalized counts was used to perform gene set enrichment analysis in GSEA v4.3.2 using the GO biological processes library. Pathways with a nominal p-val<0.05 and FDR q-val < 0.25 was used to establish significance. The matrix of normalized counts was also used to perform ssGSEA in GenePattern software using default parameters and GO library [3] and MCP-counter analysis [3] in RStudio v2024.04.01.

**Human data analysis.** *Data acquisition.* Pancreatic gene expression (n=328), whole genome sequencing (n=866, dbGaP accession number phs000424.v8), and phenotypic, epidemiological, and clinical data (n=980) were obtained from the GTEx project (https://gtexportal.org/home/)[4]. Overall, 305 individuals with the three layers of information (gene expression, genotype, and sociodemographic/epidemiological information) were considered. The studies involving human participants were reviewed and approved by GTEx data (DAR: 81372-7). The patients/participants provided their written informed consent to participate in this study.

*Imputation of the CTRB2 exon 6 deletion.* We focused on 243 individuals of European descent. To assess the *CTRB2* exon 6 deletion genotype in region chr16: 72738615-77739199 (hg38), we first filtered out loci with missing genotypes in that region and we imputed the deletion using IMPUTE2 [5], considering as reference the genotypes of 438 individuals from 1000G [6]. We then compared the imputed genotype with *CTRB2* exon 6 deletion transcriptome data determined via exon 6–7, exon 5–6, and exon 5–7 junction read counts and selected for further analysis subjects with imputed "deletion" genotype and "high exon 5-7 junction read counts and low exon 5-6 and exon 6-7 junction read counts". Eight subjects were excluded, resulting in a total of 235 subjects.

We then selected subjects overlapping with those identified by Jermusyk et al (n=174) (6). This step implies excluding individuals who were homozygous for the C allele at SNP rs1808427 (6), a proxy for the *CTRB1/CTRB2* exon 1 inversion allele variant which has been associated with chronic pancreatitis risk. Overall, a total of 159 individuals were considered (140 *CTRB2*$^{+/+}$, 17 *CTRB2*$^{+/\Delta}$, and 2 *CTRB2*$^{\Delta/\Delta}$).

*Differential gene expression analysis.* After this rigorous case identification, normalization and DGE analysis of homozygous and wild type individuals were conducted using the DESeq2 R package.

**Statistical analyses.** For comparisons showing a normal distribution of data, the two-tailed Student's t-test was used to calculate statistical significance. For comparison of data not showing a normal distribution, the two-tailed Mann–Whitney U-test was used to calculate statistical significance. All group data are represented by the mean and error bars are the standard deviation (SD). All statistical analyses were performed with GraphPad Prism v8. The statistical test used in each experiment is indicated in the Figure legends or in the Source Data file.

### SUPPLEMENTAL TABLES

**Supplemental Table 1. Top 20 genes expressed in the human and mouse pancreas***

| Human | base Mean expression (xK) | Mouse | base Mean expression (xK) |
|---|---|---|---|
| *PRSS2* | 3301 | *Ctrb1* | 589 |
| *CPA1* | 3161 | *Pnlip* | 391 |
| *PRSS1* | 2898 | *Prss2* | 321 |
| *PNLIP* | 2429 | *Cela1* | 313 |
| *GP2* | 1950 | *Prss3b* | 302 |
| *CPB1* | 1780 | *Rnase1* | 249 |
| *REG1A* | 1662 | *Cpb1* | 221 |
| *CEL* | 1314 | *Cpa1* | 212 |
| *CELA3A* | 1285 | *Cela3b* | 195 |
| *CLPS* | 1106 | *Clps* | 174 |
| *CTRB2* | 805 | *Ctrl* | 152 |
| *CTRB1* | 718 | *Try5* | 138 |
| *MT-ND4* | 691 | *Try4* | 135 |
| *CELA3B* | 615 | *Pnliprp1* | 119 |
| *CPA2* | 604 | *Cel* | 108 |
| *REG1B* | 580 | *Zg16* | 90 |
| *MT-CO1* | 577 | *Sycn* | 64 |
| *CTRC* | 502 | *Reg1* | 63 |
| *PLA2G1B* | 449 | *Cpa2* | 50 |
| *MT-CYB* | 387 | *Gp2* | 38 |

* Data derived from RNA-Seq genome wide transcriptomic analysis. The same color code is used to facilitate comparison of expression of orthologous genes.

**Supplemental Table 3. Antibodies used in this study**

| Antigen | Reference | Dilution |
|---|---|---|
| CTRB1/2 | sc-398721, Santa Cruz | IHC 1:500, IF 1:200 , WB 1:1000 |
| ATF4 | 10835-1-AP, Proteintech | IHC 1:400, WB 1:500 |
| AGR2 | HPA007912, Atlas Ab | IHC and IF 1:400, WB 1:1000 |
| BiP | ab32618, Abcam | IHC 1:200, WB 1:1000 |
| BiP | ab21685, Abcam | IF 1:200 |
| XBP-1s | E9V3E, Cell Signaling | IHC 1:100, WB 1:1000 |
| Phosphor-H2AX | 07-164, Millipore | IHC1:500 |
| P21 | HUGO291, CNIO Monoclonal Antibodies Unit | IHC1:2 |
| AMY2B | A8045, ABClonal | IF 1:400, WB 1:1000 |
| Calreticulin | ab92516, Abcam | IF 1:200 |
| Cleaved Caspase 3 | 9661, Cell Signaling | ICH 1:300 |
| CD45 | D3F8Q, Cell Signaling | IHC 1:200 |
| P53 | AM(POE316), CNIO Monoclonal Antibodies Unit | IHC 1:1 |
| CHOP | 15204-1-AP, Proteintech | IHC 1:200, WB 1:500 |
| p-eIF2a | 9721, Cell Signaling | WB 1:1000 |
| eIF2a | 9722, Cell Signaling | WB 1:1000 |
| Histone 3 | ab1791, Abcam | WB 1:1000 |

**Supplemental Table 4. Oligonucleotide primers used for RT-qPCR in this study**

| | Forward | Reverse |
|---|---|---|
| *Ctrb1_exon6* | ATTCCCTCTATCCCTGCAGC | ATGCAGGAAGAGACGCCAC |
| *Ctrb1_exon4-5* | ACTGCGGGGTCAAGACAACC | ATCGTCAACAGTGGGCAGAC |
| *Amy2b* | TGGCGTCAAATCAGGAACATG | AAAGTGGCTGACAAAGCCCAG |
| *Cpa1* | TACACCCACAAAACGAATCGC | GCCACGGTAAGTTTCTGAGCA |
| *BiP* | TGTGTGTGAGACCAGAACCG | AACACACCGACGCAGGAATA |
| *Chop* | TGTTGAAGATGAGCGGGTGG | CACGTGGACCAGGTTCTCTC |
| *Xbp1s* | AAGAACACGCTTGGGAATGG | CTGCACCTGCTGCGGAC |
| *Xbp1t* | GACAGAGAGTCAAACTAACGTGG | GTCCAGCAGGCAAGAAGGT |
| *Hprt* | GGCCAGACTTTGTTGGATTTG | TGCGCTATCTTAGGCTTTGT |
| *Cd45* | TTACCTGCTCGCACCACTG | AGCAGCGTGGATAACACACC |
| *45S* | GAGCTGGTGGTGGCGCTCC | CTTGCCCCTCCTTCTCT |
| *Mature 18S* | GATGGTAGTCGCCGTGCC | GCCTGCTGCCTTCCTTGG |
| *Mature 5.8S* | ACTCGGCTCGTGCGTC | CCGACGCTCAGACAGG |
| *Mature 28S* | GACGCGCATGAATGGA | TGTGGTTTCGCTGGATAGTAGGT |
| *5.8S 5' Junction* | TACGACTCTTAGCGGTGGATCA | TCACATTAATTCTCGCAGCTAGCT |

### SUPPLEMENTAL FIGURES

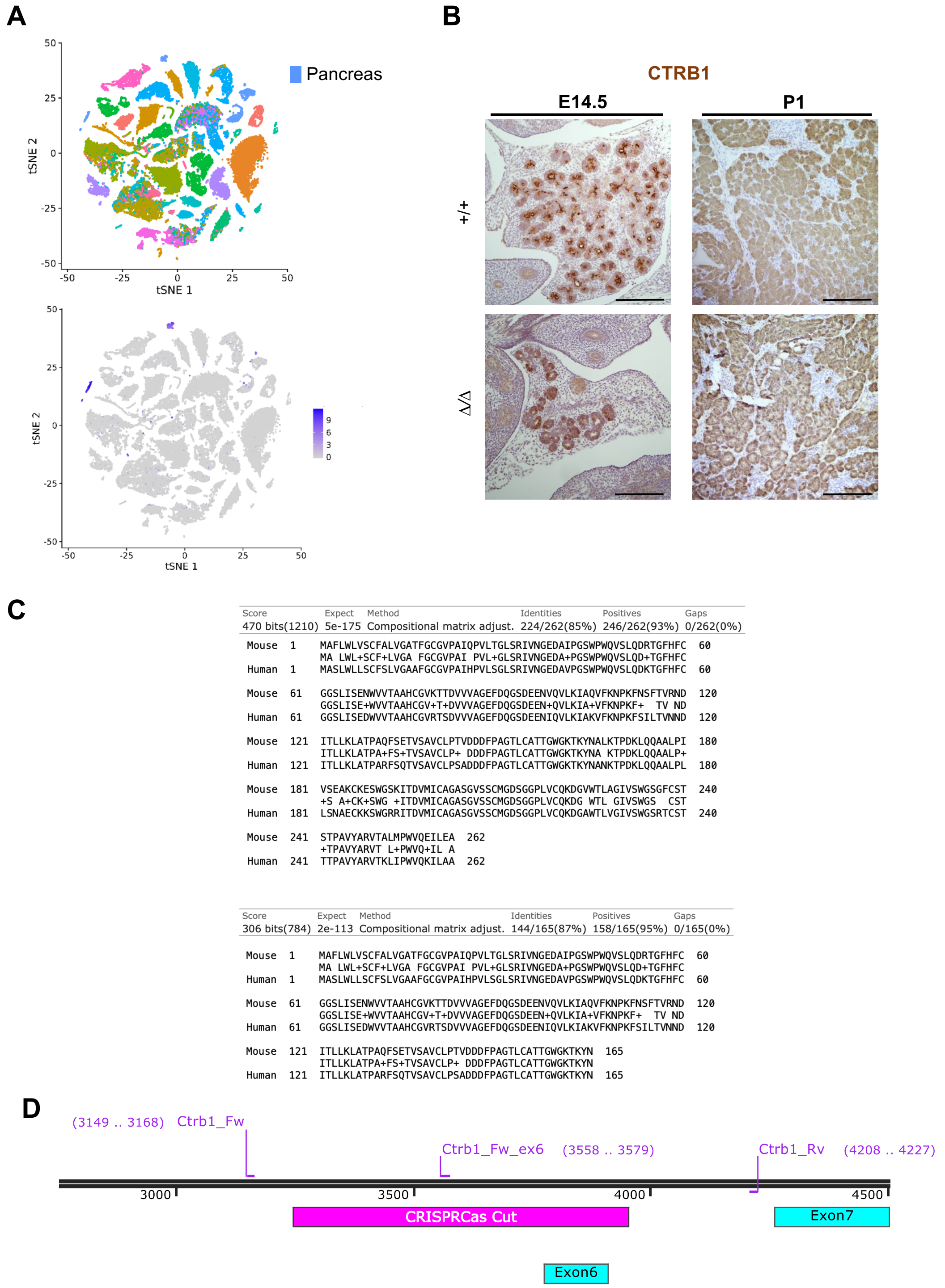


**Supplemental Figure 1**. ***Ctrb1* is expressed exclusively in the pancreas, starting at embryonic stages. Mouse CTRB1 and human CTRB2 display a high sequence identity**. **A**. tSNE plot showing that *Ctrb1* is exclusively expressed in the pancreas (data from the Tabula Muris Consortium). Scale bar units are ln(1+CMP). **B**. CTRB1 is expressed in e14.5 and postnatal day 1 pancreata. Scale bar, 100µm. **C**. Mouse CTRB1 and human CTRB2 share a 93% amino acid homology. The human and murine truncated proteins display a 95% homology. **D**. Schematic representation of the mouse genotyping strategy.

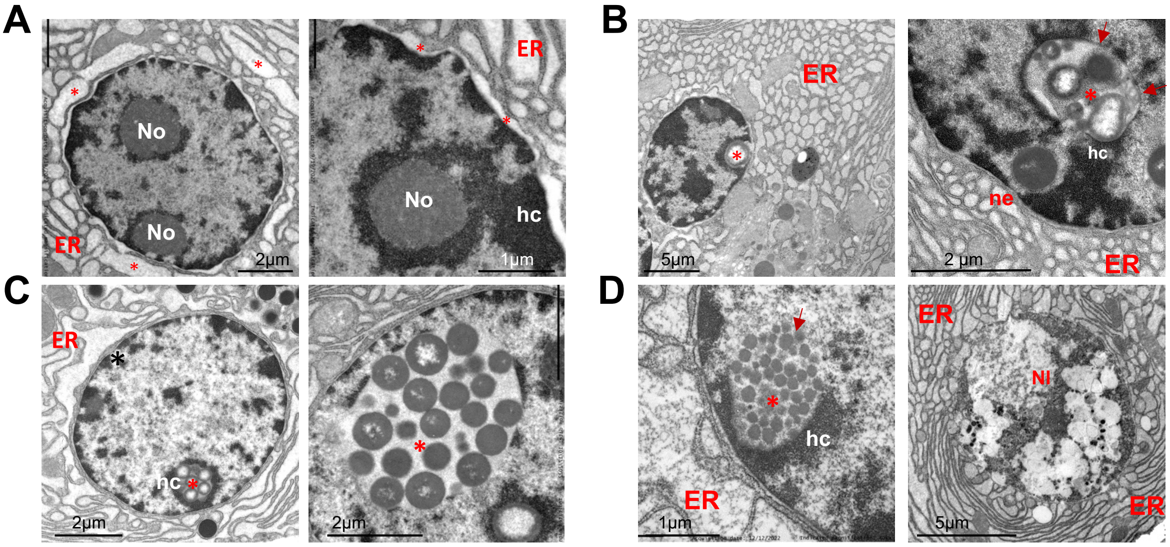


**Supplemental Figure 2. Nuclear alterations in acinar cells from *Ctrb1*$^{\Delta/\Delta}$ mice. A.** Dilation of the perinuclear and cytoplasmic ER cisternae associated with the accumulation of an amorphous, flocculent, secretory product (*). Nucleolus (No). **B.** Intranuclear inclusions are formed at the nuclear periphery (*), likely by budding/invagination or rupture of the inner nuclear membrane (red arrows). They tend to be partially covered by peripheral heterochromatin (hc). Nuclear envelope (ne). **C.** ER-stressed acinar cells frequently contain intranuclear inclusions (asterisks) with one or more secretory granules, solid or with an electron-lucent core, representing true nuclear compartments. Secretory granules are commonly embedded in an amorphous low-density (non-condensed) matrix (fluid phase), possibly derived from the perinuclear cistern. In general, there is a clear interface between the intranuclear inclusion and the nucleoplasm. **D.** Degenerative changes in acinar cells associate with rupture of the lining membrane of the nuclear inclusions (red arrow); massive release of secretory products into the nucleoplasm likely results in "nucleolysis" (nl) and cell death.

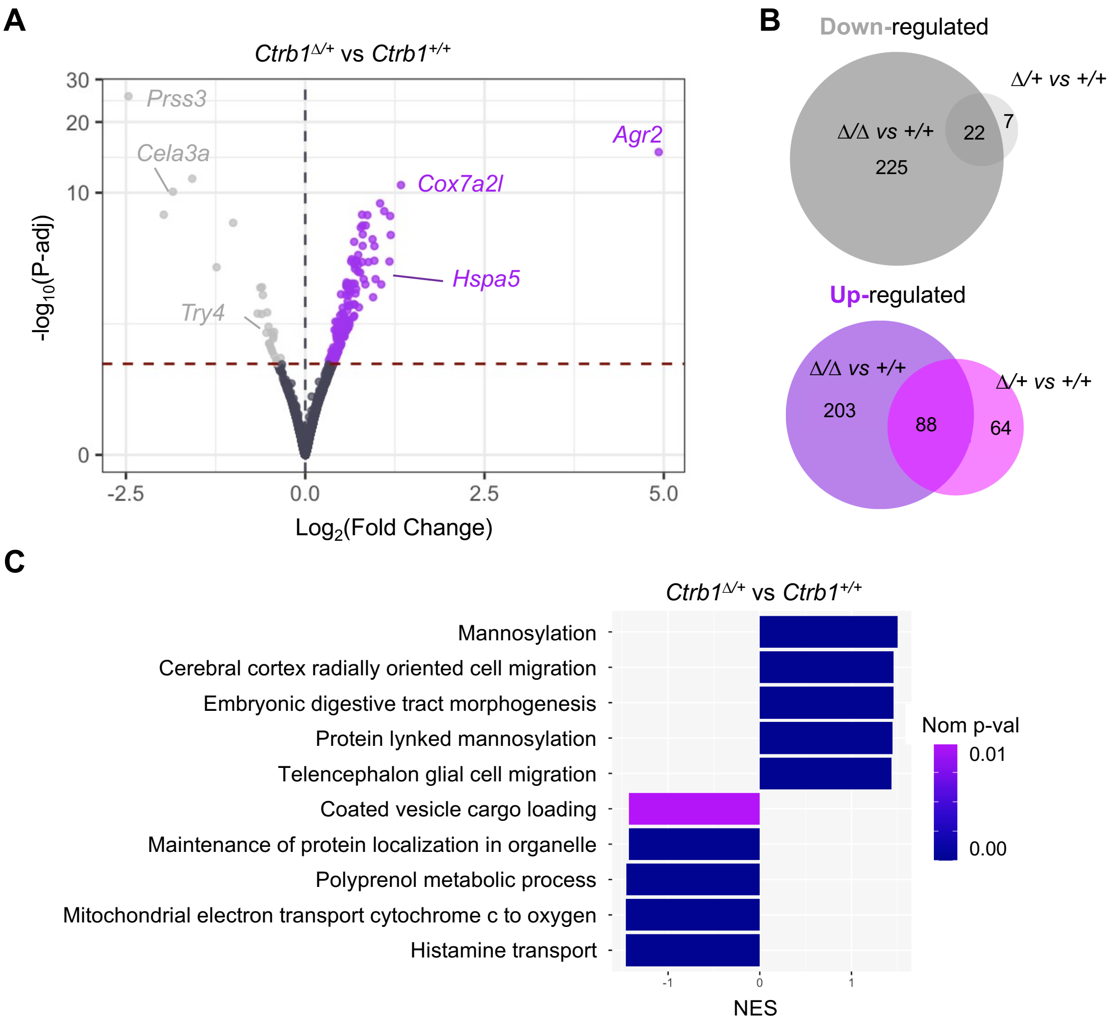


**Supplemental Figure 3. Transcriptomic analysis of *Ctrb1*$^{+/\Delta}$ *vs Ctrb1*$^{+/+}$ pancreata at 3 months. A.** Volcano plot showing differentially expressed genes (P-adj<0.05) in *Ctrb1*$^{+/\Delta}$ *vs Ctrb1*$^{+/+}$ pancreata. Up-regulated genes (n=152) are labeled in purple and down-regulated genes (n=29) in light gray. Dashed red line represents the p-adj=0.05 cutoff. Highlighted up-regulated genes are examples of ER stress-related genes and highlighted down-regulated genes correspond to digestive enzymes**. B.** Venn diagram showing the overlap between the differentially expressed genes in the *Ctrb1*$^{\Delta/\Delta}$ vs *Ctrb1*$^{+/+}$ and *Ctrb1*$^{+/\Delta}$ vs *Ctrb1*$^{+/+}$ comparisons: 88 up-regulated and 22 down-regulated genes (P-adj<0.05). **C**. Top 5 up-regulated and down-regulated pathways from the GSEA Gene Ontology gene set.

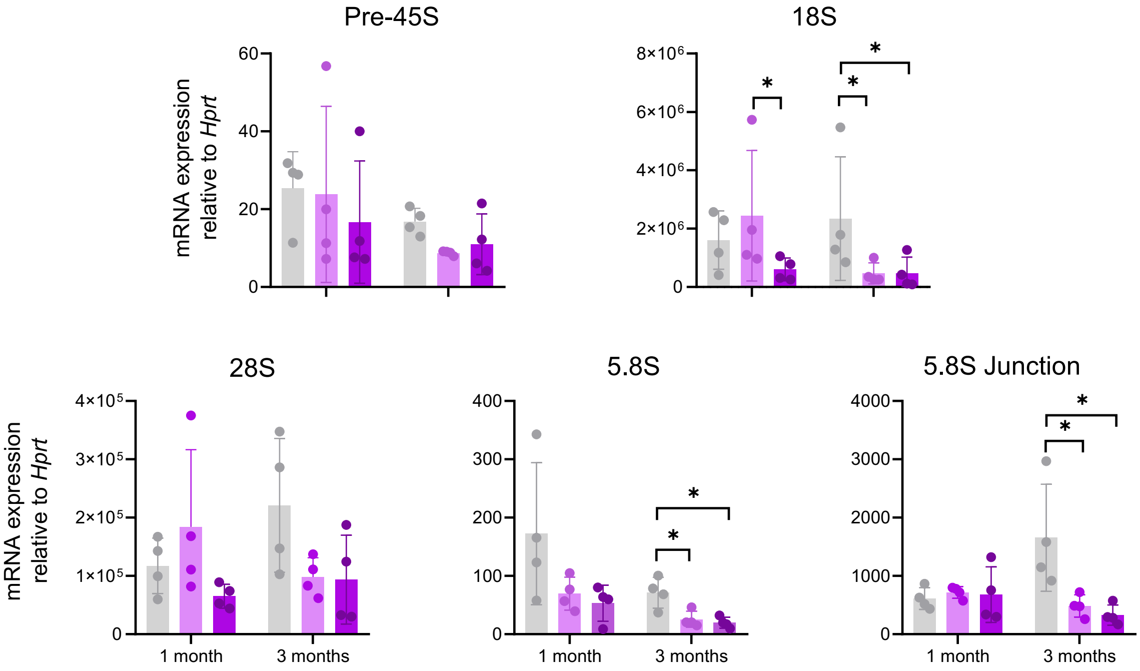


**Supplemental Figure 4. Reduced expression of rRNA transcripts in the pancreas of 3 month-old *Ctrb1*$^{\Delta/\Delta}$ mice.** Pre-45 rRNA levels are not significantly different in 1- and 3-month-old mice, regardless of the genotype. Reduced levels of 18S rRNA in *Ctrb1*$^{\Delta/\Delta}$ pancreata. 28S rRNA levels are not significantly different in 1- or 3-month old pancreata. Significantly lower levels of 5.8S and 5.8S junction amplicons in 3-month old *Ctrb1*$^{\Delta/\Delta}$ and *Ctrb1*$^{\Delta/+}$ pancreata (n=4mice/group and timepoint).

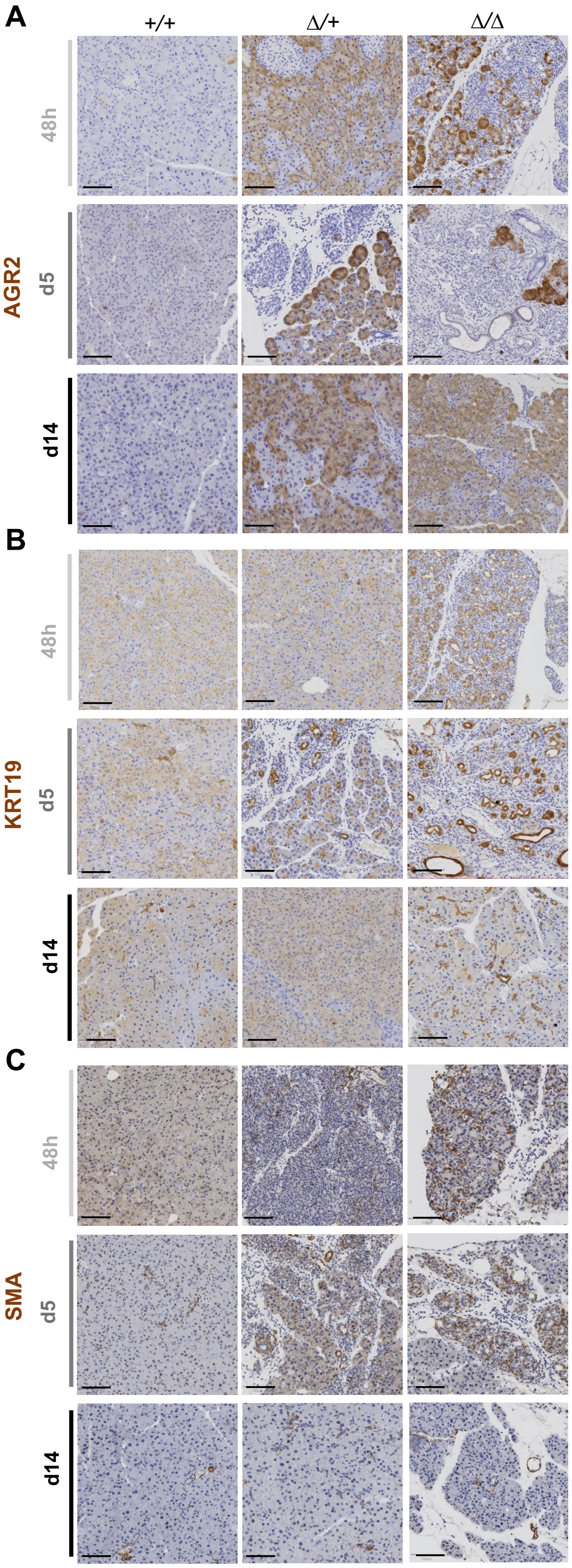


**Supplemental Figure 5. Immunohistochemical analysis of selected markers upon caerulein-induced pancreatitis. A.** AGR2 is expressed at high levels in mice carrying one or two *Ctrb1*$^{\Delta exon6}$ alleles. AGR2 expression is reduced in areas of ADM**. B** and **C.** Increased KRT19 and SMA expression in the pancreas of *Ctrb1*$^{\Delta/\Delta}$ and *Ctrb1*$^{+/\Delta}$ mice at days 5 and 14. Representative images from (n≥4 mice/group). Scale bar, 100μm.

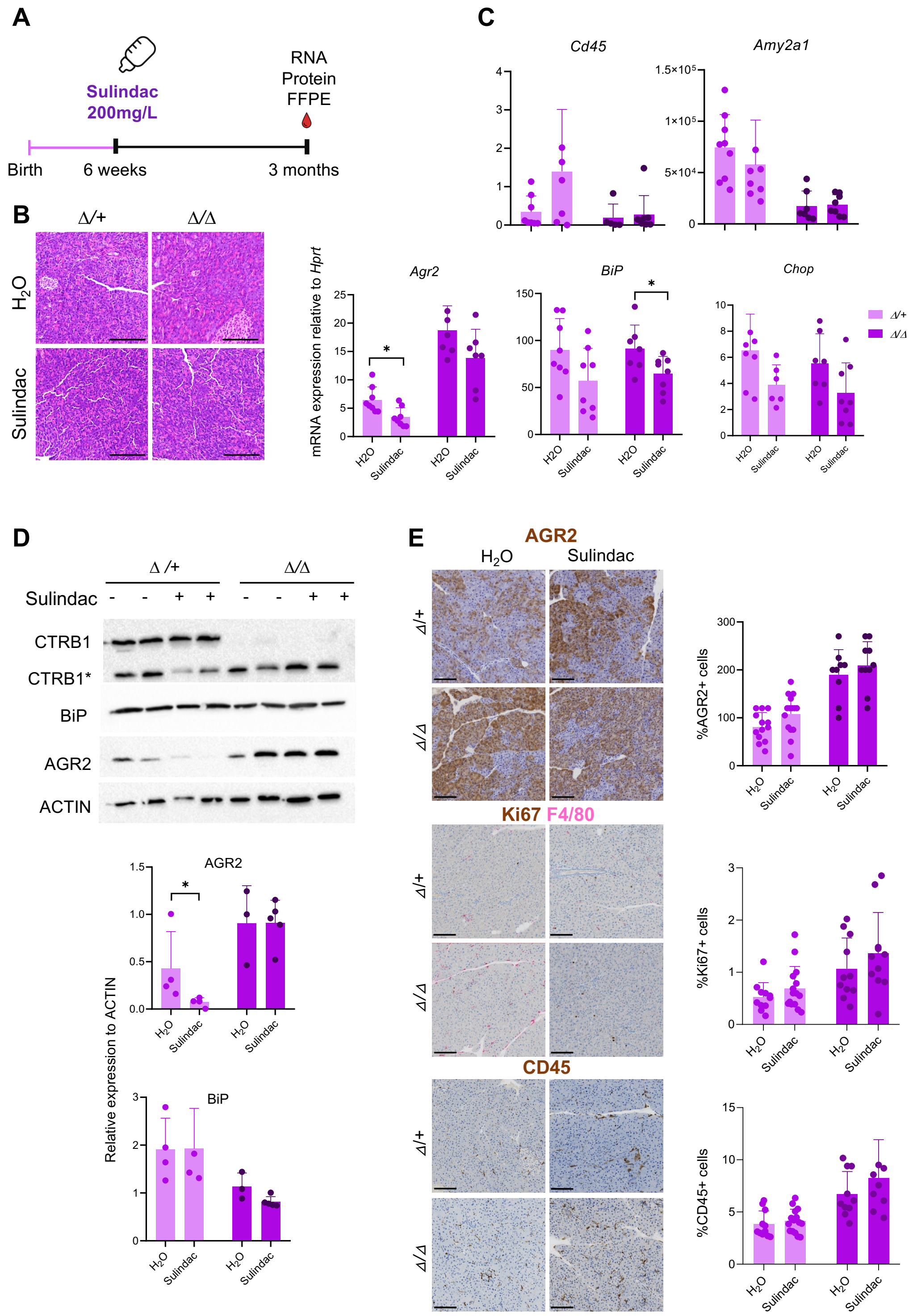


**Supplemental Figure 6. Sulindac administration to *Ctrb1*$^{\Delta/\Delta}$ mice results in a modest reduction of expression of ER stress markers in acinar cells. A.** Schematic representation of Sulindac administration: Sulindac was administered in drinking water starting at 6 weeks age until 12 weeks age. **B**. *Ctrb1*$^{\Delta/\Delta}$ and *Ctrb1*$^{+/\Delta}$ pancreata from control and Sulindac-treated mice display a normal histology. Scale bar, 100μm. **C**. RT-qPCR expression analysis of acinar (*Amy2a*), ER stress-related (*Agr2, Bip* and *Chop*), and inflammatory (*Cd45*) genes (n$\geq$6 mice/group). **D**. Western blot analysis shows that Sulindac administration results in a modest decrease of AGR2 expression in *Ctrb1*$^{+/\Delta}$ pancreata. **E**. Immunohistochemical analysis reveals similar abundance of CD45+ and Ki67+ cells in Sulindac-treated mice (n$\geq$9 mice/group). Scale bar, 100μm.
